# Differences in the free energies between the excited states of A*β*40 and A*β*42 monomers encode their distinct aggregation propensities

**DOI:** 10.1101/2020.02.09.940676

**Authors:** Debayan Chakraborty, John E. Straub, D. Thirumalai

**Author notes:** D.C., J.E.S. and D.T. designed research; D.C. performed research; D.C. and D.T. analyzed data; and D.C. J.E.S. and D.T. wrote the manuscript.

## Abstract

The early events in the aggregation of the intrinsically disordered peptide, A*β*, involve transitions from the disordered lowest free energy ground state to assembly-competent states. Are the fingerprints of order found in the amyloid fibrils encoded in the conformations that the monomers access at equilibrium? If so, could the enhanced aggregation rate of A*β*42 compared to A*β*40 be rationalized from the sparsely populated high free energy states of the monomers? Here, we answer these questions in the affirmative using coarse-grained simulations of the SOP-IDP model of A*β*40 and A*β*42. Although both the peptides have practically identical ensemble-averaged properties, characteristic of random coils (RCs), the conformational ensembles of the two monomers exhibit sequence-specific heterogeneity. Hierarchical clustering of conformations reveals that both the peptides populate high free energy aggregation-prone (*N*\*) states, which resemble the monomers in the fibril structure. The free energy gap between the ground (RC) and the *N*\* states of A*β*42 peptide is smaller than for A*β*40. By relating the populations of excited states of the two peptides to the fibril formation time scales using an empirical formula, we explain nearly quantitatively the faster aggregation rate of A*β*42 relative to A*β*40. The *N*\* concept accounts for fibril polymorphs, leading to the prediction that the less stable *N*\* state of A*β*42, encoding for the U-bend fibril, should form earlier than the structure with the S-bend topology, which is in accord with the Ostwald’s rule rationalizing crystal polymorph formation.

**Significance Statement:** Alzheimer’s disease (AD), a rampant neurodegenerative disorder, is caused by the accumulation of pathological aggregates, primarily composed of the two isoforms A*β*40 and A*β*42. Experiments have shown that A*β*42 is more aggregation-prone compared to A*β*40. However, the molecular origin of this apparent anomaly remains elusive. Here, we provide a microscopic basis for the different aggregation rates in terms of the distinct populations of high free energy excited fibril-like states (N*) that are encoded in the monomer spectra. The N* theory explains the emergence of fibril polymorphs, and predicts the relative kinetic stabilities of A*β*42 fibrils using Ostwald’s rule of stages. Our work shows that sequence-specific conformational heterogeneity of the monomer ensembles provides important cues for understanding protein aggregation.

---

**A**lzheimer’s disease (AD), a prevalent neurodegenerative disorder affecting a large proportion of the world population, is associated with the gradual accumulation of insoluble plaques in various parts of the brain, and subsequent break-down of the central nervous system (1–5). The fibrillar structures of the A*β* peptides, like other amyloid aggregates, have a characteristic cross-*β* architecture (6–8). The fibrils form as a result of aggregation of the 39-43 residue long amyloid-A*β* peptides. *In vivo*, A*β* is produced by the proteolytic cleavage of the amyloid precursor protein (APP) by *γ*-secretases, with A*β*40 and A*β*42 being the major products (9, 10). The fibrils exhibit a high degree of polymorphism, even though globally they all have the characteristic *β*-sheet architecture. The link between polymorphic structures, which could depend on the way they are generated, and the extent of neurotoxicity is a topic of continuing interest (1, 11). In addition to AD, several other neurodegenerative diseases, such as Parkinson’s and Huntington’s disease, are also the result of abnormal *in vivo* protein aggregation. Various studies (12, 13) have proposed that these diseases could share several common themes with AD, including similar fibril morphologies, mechanism of fibril formation, and the association of toxicity with oligomers. To obtain a more quantitative understanding of these general principles, it is crucial to understand the key steps involved in amyloid aggregation at the molecular level.

The cascade of events that describe the conversion of monomers to fibrils is complex (14–16). At each step, it is likely that an ensemble of highly heterogeneous conformations are populated. Because of the transient population of species that form during the early stages (before a critical size nucleus) it is difficult to describe their structural details using experiments alone. A complete understanding requires describing the conformational changes that occur during the aggregation process starting from the monomer. Towards this end, conformational dynamics of A*β* monomers have been extensively studied, both using experiments (17–19), and computer simulations (15, 20–25). The structural details of the predicted ensembles vary widely among the different studies. Some of the differences in the experiments could be due to differing external conditions, such as variations in pH, temperature or salt concentration. Molecular dynamics simulations reveal that the populations of different secondary and tertiary structural elements are strongly dependent on the details of the force field and sampling strategies (26–29). However, based on recent NMR (17, 18) and FRET experiments (30), a general consensus seems to be emerging that A*β* monomers behave like many other Intrinsically Disordered Proteins (IDPs) that are highly disordered, adopting random coil (RC) structures at room temperature, and neutral pH. The structures are devoid of any persistent secondary structures (*α*-helices and *β*-sheets). In other words, it appears that the propensity to form amyloid fibrils cannot be discerned when only the average properties of the monomer ensemble are examined.

The natural question is if there are any connections between the conformational heterogeneity of the monomer ensemble in solution, and the eventual structures adopted in the fibril state. More precisely, at what stage of A*β* aggregation does one observe a transition from the RC conformation to structures with considerable *β* content? In a series of papers, we (31–33) showed that the structures that have a high propensity to aggregate are encoded as excitations in the monomer free energy spectrum. Subsequent studies (34–36) have confirmed our findings. More importantly, such conformations have some of the structural features of the monomers in the fibril. We first elucidated this concept using atomically detailed simulations of a 26 residue fragment of the A*β* peptide using molecular dynamics simulations (31). Subsequently, we illustrated the consequences quantitatively using lattice models (32), for which precise computations could be carried out by exploring the entire sequence space. Simulations based on the lattice models also showed empirically that the time scales of fibril formation (*τ*_*fib*_), could be linked to the population (*p*_*N*\*_) of the aggregation-prone species (referred to from now on as *N*\* states) (32, 36). The structures in the *N*\* ensembles (there is more than one) are difficult to characterize using standard experimental techniques because they are high free energy states that are only sparsely populated. In this context computations are useful.

For A*β*40 and A*β*42, the two most prevalent isoforms *in vivo*, the question whether there is a link between the underlying heterogeneity (and by inference, *p*_*N*\*_) of the monomeric ensembles, and their aggregation propensities is pertinent. This is because A*β*42 aggregates faster, and is the major constituent of amyloid plaques (37–39), despite being present in relatively lower abundance than A*β*40 (40). Several studies have attributed the elevated pathogenicity of A*β*42 to the pronounced structuring near the C-terminus, which is mediated by the two extra residues (ILE41 and ALA42) (41–43). Recent A*β*42 fibril structures determined using solution-state NMR (44, 45), and Cryo-EM (46) seem to corroborate this viewpoint. Previous work based on all-atom studies (29, 30, 42, 47, 48) simulations using coarse-grained models (49–51) further predict that the two isoforms could be associated with different populations of the metastable states, and allude to a sequence-specific conformational heterogeneity that could be a key determinant of the contrasting aggregation propensities.

Here, we report results from simulations using a modified version of a recently developed highly accurate SOP-IDP coarse-grained (CG) model for (IDPs) (52) to characterize the conformational ensembles of A*β*40 and A*β*42. The major purpose is to illustrate the theory that the differences in the extent of population of the the sparsely populated *N*\* states in the monomeric forms of the A*β* peptides quantitatively explains the relative fibril elongation rates of A*β*42 and A*β*40. We show that despite having practically identical ensemble-averaged values of several experimental observables under physiological conditions, the underlying conformational heterogeneities of the two peptides are distinct. In particular, the termini display different extent of residual structuring in the two peptides. Most importantly, A*β*42 has a higher population (*p*_*N*\*_) of aggregation-prone (excited) states that display many of the characteristic structural features found in fibrils. The difference in the equilibrium populations of *N*\* states between A*β*42 compared to A*β*40 when linked to kinetics rationalizes the nearly one order of magnitude faster fibril elongation rate of A*β*42 relative to A*β*40. Our study, therefore, provides a crucial link between aggregation propensity and sequence-specific conformational heterogeneity in the monomer. In other words, the spectrum of excited, but sparsely populated *N*\* states sampled at equilibrium, is a harbinger of protein aggregation.

## Results

### Sizes and polymeric features of A*β* peptides are nearly identical

The distributions of the radius of gyration, *R*_*g*_, end-to-end distance, *R*_*ee*_, both of which describe the global dimensions of the A*β* monomers, shown in Figure 1, are not significantly different. The small differences are likely due the longer length of A*β*42. The ensemble averages of these observables for the two sequences are nearly identical (Table 1). Our estimates for ⟨*R*_*g*_⟩ and ⟨*R*_*ee*_⟩ are in good agreement with the values reported in two recent studies (28, 30). The distributions of the scaled end-to-end distance, 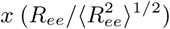, shown in Fig. 1(c), for both the A*β* monomers deviate substantially from the well-established theoretical results for a Gaussian chain, and a self-avoiding random walk (SAW) (see SI for further details). The departure from standard homo-polymer theories is the first hint that sequence-specific effects as well as the inherent polyampholyte-like characteristic of the A*β* peptides, that are masked in moments of the distribution functions, need to be accounted for when describing their conformational ensembles.

**Table 1.**
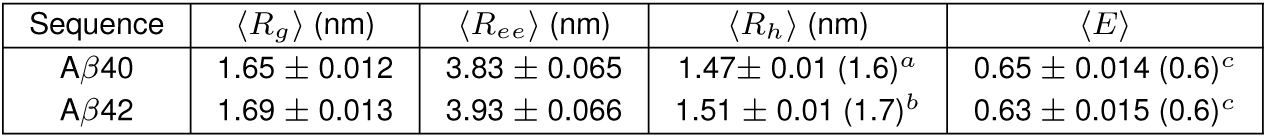
The ensemble-averaged values of different observables for A*β*40 and A*β*42. The errors are reported in terms of standard deviations estimated using block-averaging. ^*a*^ The experimental value of *R*_*h*_ is from FCS experiments of Wennmalm *et al.* (54). ^*b*^ The experimental value of *R*_*h*_ is from the diffusion NMR experiments of Vendruscolo and coworkers (53). ^*c*^ The FRET efficiency values are from Meng *et al* (30). The numbers in parenthesis are taken from the quoted experiments.

**Fig. 1.**
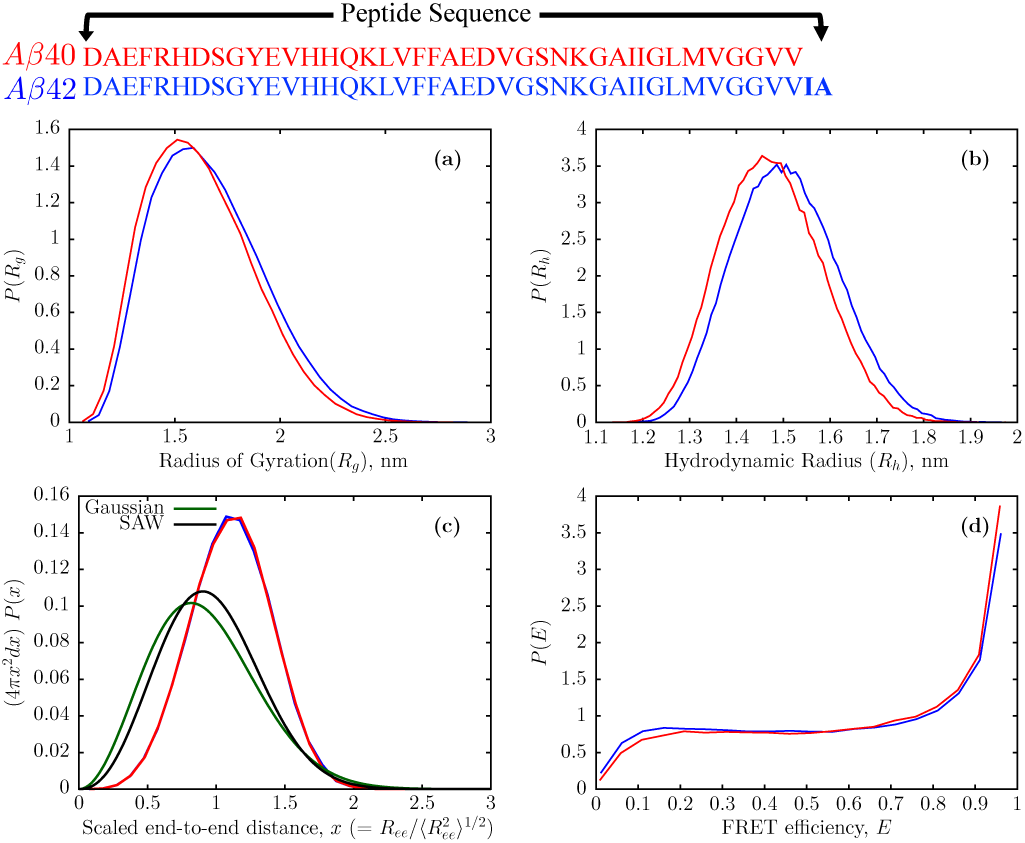
The sequences of A*β*40 and A*β*42 using one letter code is displayed on top of the figure. Ensemble characteristics of A*β*40 and A*β*42 monomers depicted as red, and blue curves, respectively. Distributions of the radius of gyration (a), hydrodynamic radius (b), end-to-end distance (*R*_*ee*_) scaled by 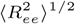. To compare with the results for a Gaussian chain, and a self-avoiding walk (SAW), we show 4*πx*^2^*P* (*x*) where 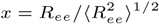, and FRET efficiencies (d). The corresponding mean values are given in Table 1.

Besides *R*_*g*_ and *R*_*ee*_, the hydrodynamic radius *R*_*h*_ is another measure of a polymer size, which is routinely estimated using dynamic light scattering (DLS), NMR, and FCS experiments (53, 54). The distributions of *R*_*h*_ for the A*β*40 and A*β*42 also overlap to a large extent (Figure 1(b)), and yield similar ensemble-averages (Table 1). For both the sequences, the ⟨*R*_*h*_⟩ values are in excellent agreement with (1.5-1.7 nm) NMR (53) and FCS experiments (54). The ratio *R*_*h*_*/R*_*g*_ for both A*β*40 and A*β*42 is ≈ 0.9, which implies that neither peptide behaves as a Gaussian chain, for which the ratios should be 0.640 and 0.665, respectively (55).

In a recent study (30), Meng *et al.* reported the global dimensions and conformational dynamics of A*β* monomers using single-molecule experiments, thus complementing previous findings from NMR and CD experiments. The FRET histograms computed from our simulations, using Eq. 8, are shown in Figure 1(d). The average FRET efficiencies, ⟨*E*⟩ for both the peptides (Table 1) are in excellent agreement with the experimental estimates (≈ 0.6) at zero denaturant concentration (30). Unlike in the FRET experiment, however, we also observe a peak at ⟨*E*⟩ ≈ 1. This discrepancy is not entirely unexpected, and could be attributed to the small size of the peptides, as well as the large Förster radius (5.2 nm) of the Alexa488 and 647 dye pair, which according to Eq. 5, would result in ⟨*E*⟩ ≈ 1 for *R*_*ee*_ less than 4.0 nm. Similar observations have been made in two previous studies (28, 30). From all-atom simulations, Meng *et al.* noted that although the A*β* monomers were characterized by largely overlapping distributions for *R*_*g*_, *R*_*ee*_ and ⟨*E*⟩, the *R*_*ee*_ distribution of A*β*42 exhibited a minor peak at low end-to-end distances, corresponding to a subpopulation of conformations with long-range contacts. However, no such feature is discernible from our computed probability distributions (Figure 1). The good agreement between our simulations and experiment for the observables in Table 1 attests to the accuracy of the SOP-IDP model in predicting the values of measurable quantities.

### Few metastable states cannot represent the A*β* ensembles

For both the sequences, the inter-residue distances, *R*_*ij*_ as a function of sequence-separation, |*i* − *j*|, adheres to the Flory scaling law: *R*_*ij*_ = *R*_0_ |*i* − *j*|^*ν*^. In fact, the scaling behaviors are found to be identical (Figure 2) with the Kuhn length, *R*_0_=0.47 nm, and the scaling exponent, *ν*=0.58. The value of *ν* suggests that A*β* monomers behave as polymers in a good solvent. However, we should note that *ν* estimates from either simulations or experiments suffer from finite-size effects (56).

**Fig. 2.**
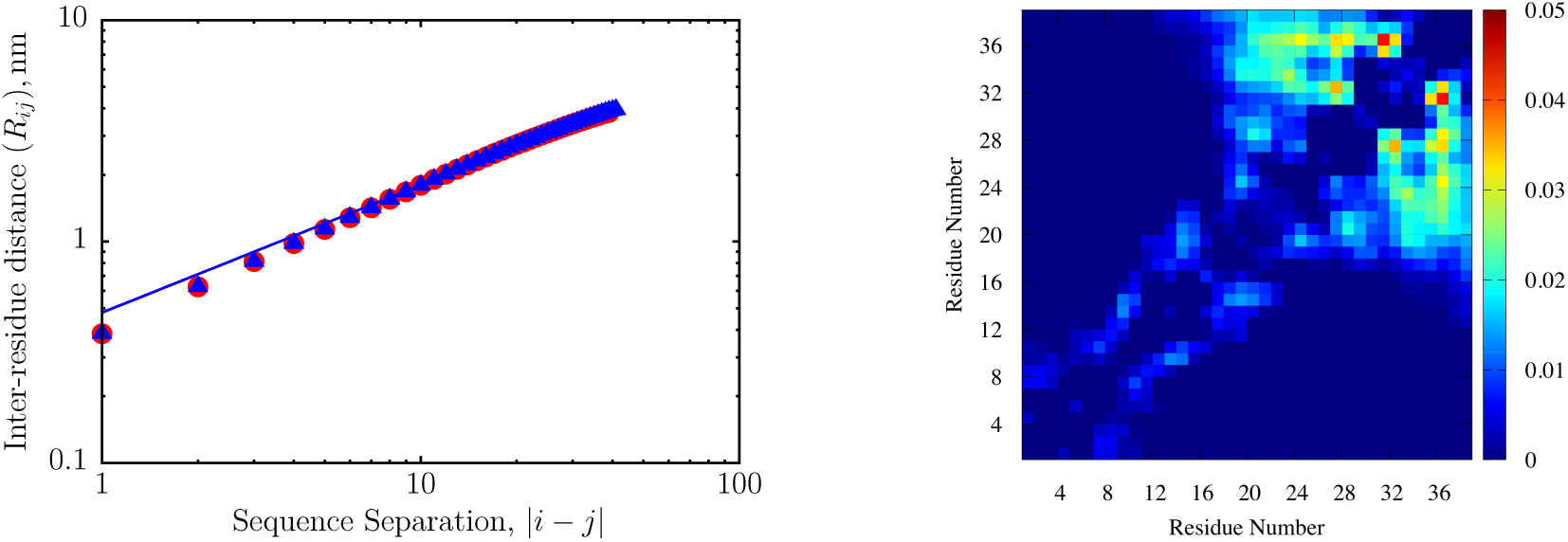
Left panel: The variation of inter-residue distance (*R*_*ij*_) with sequence separation, |*i* − *j*| for the A*β*40 and A*β*42 sequences are shown as red and blue circles, respectively. The solid lines are fits to the Flory scaling law expected for homopolymers in good solvents: *R*_*ij*_ = *R*_0_ |*i* − *j*|^*ν*^, with *R*_0_ = 0.47 nm and *ν* = 0.58 for both the peptides. Although *R*_*ij*_ scaling, an ensemble averaged quantity, is consistent with polymer theory the distribution of *R*_*e*_*e* deviates from the expected universal behavior for self-avoiding walks (see Fig. 1). Right panel: Variation in the ensemble-averaged side-chain side-chain contacts for the A*β*40 and A*β*42 shown as a difference map, color coded (scale on the right) in terms of 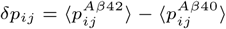. Here, 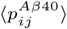 and 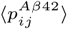 are the ensemble-averaged contact probability maps for A*β*40 and A*β*42, respectively.

The ensemble-averaged contact maps for A*β*40 and A*β*42 do not reveal discernible persistent long-range contacts (Figure S2), illustrating the disordered nature of the conformational ensembles. However, from a zoomed-in view of the difference map, *δp*_*ij*_, over a low probability range, it is apparent that a small but detectable population of substates (masked in the ensemble-averaged contact map), which have a tendency to form residual structure at the C-terminus is exclusively present in the conformational ensemble of A*β*42 (Figure 2). We describe this feature in detail, and establish possible connections to aggregation propensity in subsequent sections.

The free energy landscapes of the A*β* monomers projected onto the first and second principal components, based on the C*α*-C*α* distances, further exemplify the intrinsically disordered nature (as well as the underlying conformational heterogeneity) of the conformational ensembles (Figure S3). Consistent with previous studies on A*β*40 and 42 monomers (29, 53, 57), and other aggregation-prone peptides (58), we find that the free energy landscapes do not exhibit well-separated basins of attraction. Instead, they consist of a single featureless contour. The free energy landscapes projected onto the *R*_*g*_ and *R*_*ee*_ coordinates also exhibit similar characteristics (Figure S4) suggesting that the topographical features of the surface are largely unaffected by the particular choice of the order parameters.

Based on the ensemble-averaged contact maps, scaling behavior, as well as the nature of the underlying landscapes, we surmise that A*β* monomers do not adopt stable well-defined secondary or tertiary structures at equilibrium, but rather dynamically interconvert among a menagerie of conformations, whose relative populations can be tuned by external conditions, such as salt concentration, pH, temperature, crowding or partner proteins.

### A*β* monomers have negligible residual secondary structure

The secondary structure profiles associated with the A*β*40 and A*β*42 monomers are shown in Figure 3. Both the peptides have very low preferences for structural elements, such as *β*-strand or helices. All the residues display a very high propensity (> 97%) to form turns or irregular coils. Our observations are consistent with NMR experiments (17, 18), as well as all-atom simulations (28, 30), which predict that A*β* monomers populate mostly random-coil (RC)–like structures in equilibrium. Overall, we observe a large overlap between the turn profiles for A*β*40 and A*β*42. In particular, the elevated turn propensities near residues 7-11, as well as 23 to 27 (which includes the well-known VGSN turn) is consistent with the NMR experiment (18). Nonetheless, there are some subtle differences in the propensity of forming residual structure in some parts of the sequence, which are worth noting.

**Fig. 3.**
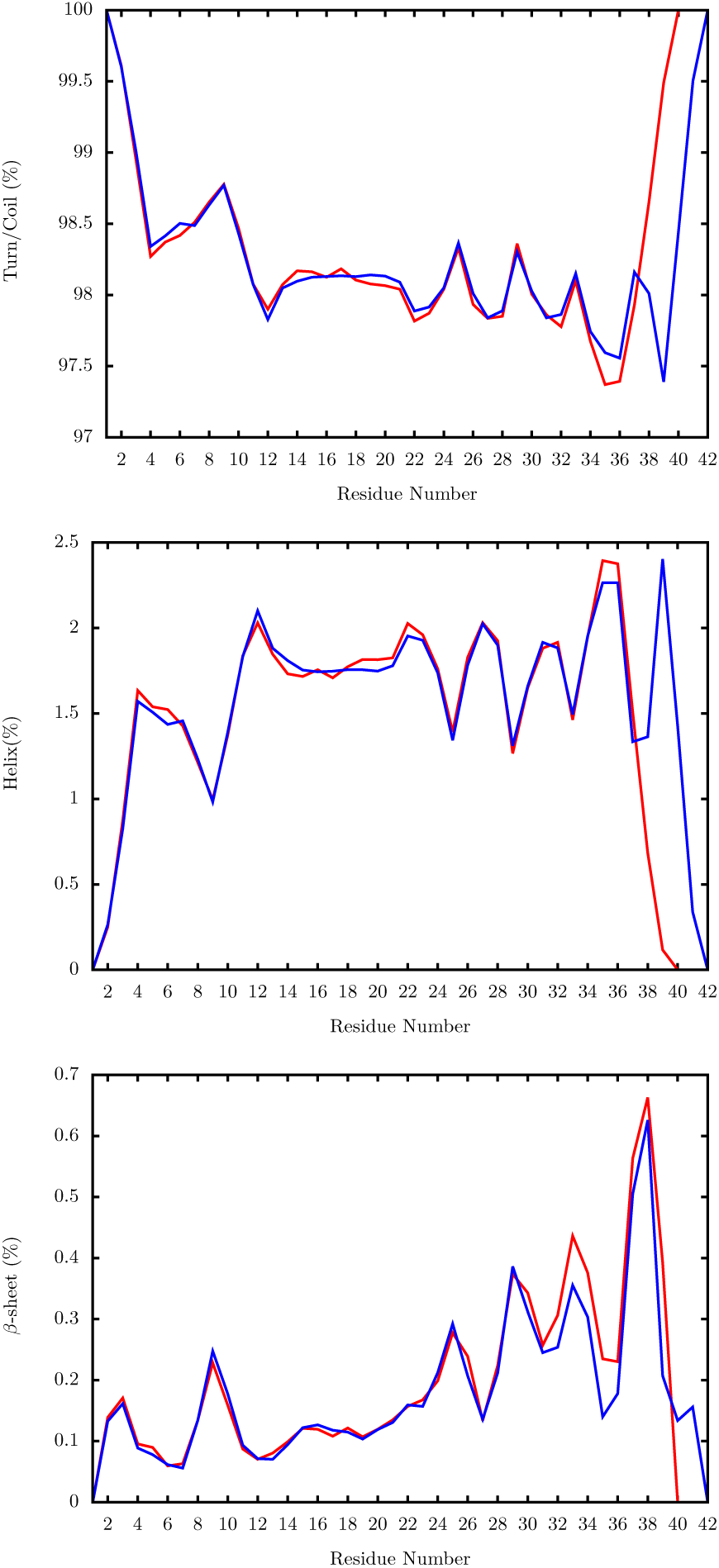
Percentages of different secondary structure elements in the conformational ensembles of A*β*40 (red), and A*β*42 (blue). Top panel: Ensemble-averaged percentages for forming turn/irregular coils. Middle panel: Residue-wise percentages for forming helices. Bottom panel: Residue-wise percentages for forming *β*-sheets. The secondary structures were assigned based on the positions of the C_*α*_ positions using the PCASSO algorithm.

A*β*40 has a more structured N-terminus (residues ≈ 4 to 6), as indicated by the marginally higher propensity to form *β*-sheets and *α*-helices in this region. In contrast, A*β*42 has a slightly more ordered C-terminus (≈ 38 to 42), with the residues in this region displaying a somewhat higher probability of forming both helices and *β*-sheets, and a pronounced bend near residues 36-37. The contrasting structural preferences at the two termini, particularly the enhanced ordering at the C-terminus of A*β*42, have also been noted in previous allatom (26–28, 42, 59), and coarse-grained simulations (49, 60). Interestingly, in both the peptides, residues near ≈ 13 to 20, which include the central hydrophobic core (CHC), show an enhanced probability to form helical domains. As argued in previous experimental (61), and simulation studies (23, 62, 63), such residual structure near the CHC could play a key role in mediating the early events in amyloid fibrillogenesis. We also find that in both the monomers, residues 29-36 tend to form secondary structures (specifically *β*-turns for A*β*42) with higher probability compared to the rest of the sequence. This feature is also consistent with previous work (18, 27, 48). It is worth emphasizing that the probabilities to adopt *β*-sheet or *α*-helix are really small relative to coil-like states. Our simulations suggest that detecting them using experiments is likely to be difficult.

### Chemical Shifts

The residue-wise chemical shifts for A*β*40 and A*β*42 are shown in Figure 4. Given the coarse-grained nature of our model, and the higher errors associated with the estimation of chemical shifts of other nuclei with the LARMOR-C_*α*_ formalism (see Methods) (64), we report only the ensemble-averaged values of the C_*α*_ chemical shifts. As illustrated in Figure 4, our predictions are in reasonable agreement with NMR experiments (17, 18). Despite only achieving qualitative agreement (Figure S5), it is nevertheless satisfying because we did not adjust any parameter to fit the experiments. The chemical shifts for the first thirty-five residues are nearly identical for both the A*β* monomers, mirroring the observed trend for the secondary structure profiles.

**Fig. 4.**
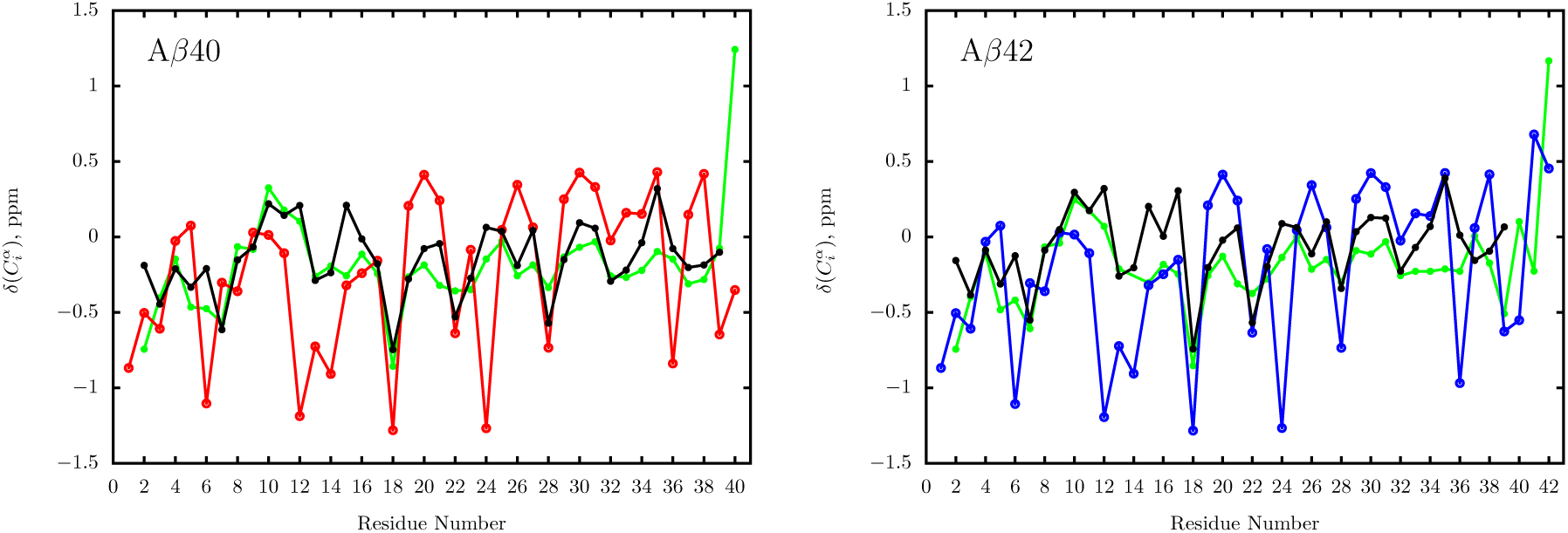
Residue-wise C_*α*_ chemical shifts for A*β*40 (left) and A*β*42 (right). The predicted shifts for A*β*40 are shown in red, and for A*β*42 are shown in blue. The chemical shifts from the experiment of Bax and coworkers are shown in light green (17), and those from Zagorski and coworkers are shown in black (18). The predicted values are in qualitative (the trends are comparable) agreement with experiment. The weighted-root-mean-squared-error (*ζ*) for A*β*40 between predicted shifts, and those measured by Bax and coworkers is 0.53. The error is smaller (0.47) relative to the measured values of Zagorski and coworkers. For A*β*42, the corresponding *ζ* values are 0.54 and 0.49.

Most residues show only minimal departures from the random coil (RC) references, further demonstrating the completely disordered nature of the ensembles, and the lack of substantially populated (metastable) structural elements. Nonetheless, the chemical shifts for some valines (Vs) and histidines (Hs) deviate from the RC values. In particular, the upshifted C_*α*_ shifts for V18 (also visible in the experimental profiles), and V12-H14 for both A*β* monomers implies that this region could adopt residual *β* structure, albeit only transiently. Similar observations have been made previously (65) using EPR spectroscopy, and all-atom simulations (34, 48). The chemical shifts for H6, V24 and V36 are also shifted upfield by ≈ 1 ppm, indicating that these residues could form fleeting *β* structures. Although similar signatures are not present in the experimental profiles shown in Figure 5, a previous study (48) does suggest that these residues could be involved in hydrogen-bonding interactions with V18, resulting in upshifted chemical shifts. Interestingly, the modest departure from the RC values for H6 and H14 hints at a possible role of these residues in mediating A*β* self-assembly (18, 66) via formation of transient *β* structures, as well as being a template for molecular recognition (67).

**Fig. 5.**
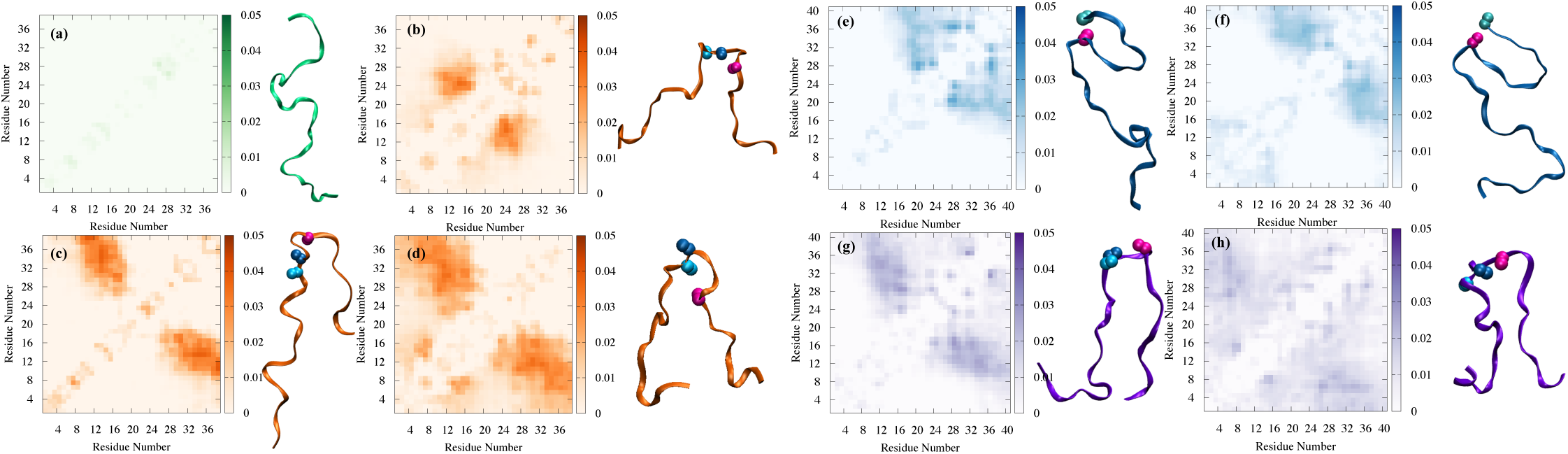
(a) The free energy ground state of the A*β*40 and A*β*42 ensembles consisting of RC-like conformations devoid of fibril-like contacts. The contact map averaged over the corresponding sub-ensemble is featureless. (b), (c) and (d) show the contact maps and representative snapshots of clusters identified for A*β*40, which consist of aggregation-prone (*N*\*) conformations. These conformations exhibit many of the structural elements found in the A*β*40 fibril. The E22, D23 and K28 residues, which form the key salt bridges, are explicitly shown. (b) Cluster 9: the VGSN turn is formed, and is stabilized by the D23-K28 salt bridge. However, the N-terminal and C-terminal segments are quite flexible, and do not form contacts. (c) Cluster 10: A hairpin like structure stabilized by the formation of a VGSN turn, and the D23-K28 salt bridge. (d) Cluster 11: Hairpin-like structure stabilized by a turn formed near the VGSN region, and the E22-K28 salt bridge. (e), (f), (g) and (h) show the contact maps and representative snapshots of clusters identified for A*β*42, which consist of conformations, some of which have structural elements found in the different A*β*42 fibril polymorphs. These structures are members of the *N*\* ensemble. In (e) and (f) the K28 and A42 residues are shown as spheres. (e) Cluster 4: S-bend type motif, exhibiting close contact between K28 and A42, similar to that found in a A*β*42 fibril structure. (f) Cluster 5: S-bend type structures, with enhanced ordering near the C-terminus, including a close contact between the K28 and A42 residues. (g) and (h) denote clusters which bear resemblance to the monomer unit of a different A*β*42 polymorph, which is a classical *β*-hairpin consisting of a U-bend. (g) Cluster 7: Hairpin-like structure consisting of a U-bend near the VGSN turn region, stabilized by the D23-K28 salt bridge (shown as spheres). (h) Cluster 6: Hairpin-like structures formed via the association of residues ≈ 25-35 and the N-terminus. The D23-K28 salt bridge (shown as spheres) stabilizes the turn region in these structures.

Given that all the measurable and computable values are virtually identical for both the peptides is it possible to explain the enhanced aggregation rate of A*β*42 relative to A*β*40 using equilibrium monomer properties? We show below that the answer lies in the differences in the conformational heterogeneities between the two peptides, which result in enhanced population of *N*\* states in A*β*42 compared to A*β*40.

### Sequence-dependent conformational heterogeneities

The ensemble-averaged properties, as well as features of the free energy landscapes, although commensurate with the prevailing consensus regarding the intrinsically disordered nature of the A*β* monomeric ensembles, do not reveal significant alloform-specific differences. Structural details of aggregation-prone conformers having extremely low equilibrium weights are masked in such averages, although subtle differences in the secondary structure profiles allude to sequence-specific conformational heterogeneity. To glean further insights into the fine structure of the free energy landscape of A*β*40 and A*β*42, we carried out hierarchical clustering of the conformational ensembles based on a robust distance metric (Figure S6). Our previous study (52) showed that this scheme provides an efficient means to quantify the contrasting conformational heterogeneities of many IDPs.

At a coarse level, the conformational ensembles of both the peptides can be partitioned into three major clusters: compact, semi-compact, and extended. Even when such simplistic classification is used, sequence-specific conformational preferences become apparent (Figure S5). For both the sequences, the equilibrium weights of the fully extended structures are relatively low, being 19.9% and 20.9% for the A*β*40 and A*β*42 sequences, respectively. Compact structures dominate the conformational ensemble of A*β*40 (53.7%), and semi-compact structures constitute the second largest cluster (26.4%). This trend is reversed for A*β*42; the population of semi-compact structures is enhanced to 41%, and those of compact structures is diminished to 38.1%. The relative weights of the different cluster families, even at such a coarse resolution, hint at evident deviations from standard polymer models, even though ensemble-averaged contact maps and scaling behavior suggest otherwise.

To ascertain the arrangement of the local structural segments we performed more refined analyses using a smaller distance cutoff for *D*_*ij*_ (see Eq. 12). This splits the dendrogram further, and divides the major cluster into additional subfamilies. A cutoff of 5.0 (see Methodology section) seems appropriate for both the A*β*40 and A*β*42 as it not only provides sufficient resolution to determine the key differences in the contact maps of the sub-clusters, but also keeps their number tractable. Using this scheme, we obtained 12 clusters for A*β*40, and 11 clusters for A*β*42 (Figure S6). The populations of the different clusters are tabulated in the SI (Table S3). In Figures 5 and 6 we show the most important cluster families that are related to the *N*\* states, and the rest are included in the supplementary information (Figures S8 and S9). The proper segregation of the various families in the two dimensional surfaces defined by *R*_*g*_ and *R*_*ee*_ implies that our clustering scheme is robust (Figure S7). The structural details of the different subfamilies are described in detailed below.

**Fig. 6.**
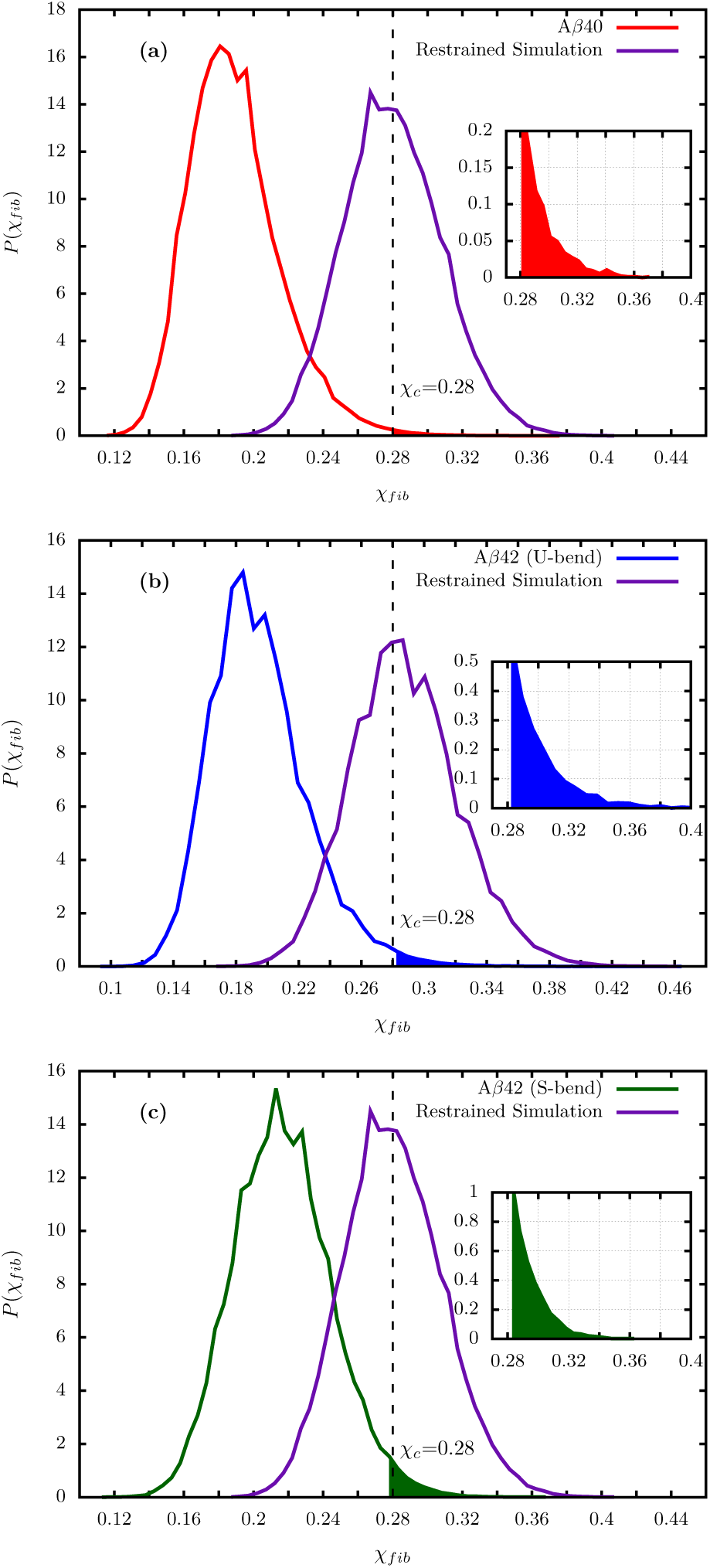
(a) Distribution of *χ*_*fib*_ for A*β*40 (shown in red). The overlap parameter is computed with respect to the monomers in the experimental fibril structure. For A*β*40, the reference state is the monomer unit from the striated (U-bend) fibril structure (76) (PDB ID: 2M4J) (76). The shaded area under the curve which corresponds to the *N*\* state is in red. For clarity an enlarged view of this section of the curve is shown in the inset. (b) Distribution of *χ*_*fib*_ for A*β*42 (shown in blue) using the monomer from the U-bend fibril structure (PDB ID: 2BEG) (72) as a reference. The shaded area under the curve (and the inset) corresponds to the population of *N*\* states with U-bend topology. (c) Same as (b) except the is calculated using the S-bend fibril structure (PDB ID: 2NAO) (44). The shaded area under the curve (and the inset) corresponds to the population of *N*_*_ states having S-bend topology. In (a), (b) and (c) the purple curves correspond to *P* (*χ*_*fib*_) computed from the restrained simulations (see SI for details) starting from the monomer present in the fibril structure. (b) A*β*40 fibril structure (PDB ID: 2M4J); (c) A*β*42 U-bend fibril structure (PDB ID: 2BEG); (d) A*β*42 S-bend fibril structure (PDB ID: 2NAO). In all cases, the center of the distribution is approximately at *χ*_*fib*_ = 0.28. This value corresponds to *χ*_*c*_, and is marked in the plots.

### A*β*40

The contact maps associated with clusters 1, 2 and 3 and 7 (Figure S8) are devoid of short or long-range contacts. Other clusters are also rich in RC-like conformations, but also contain some fingerprints in the contact map, which is suggestive of local structuring near the N-terminus. Approximately 50% of the conformations in clusters 5 and 8 show enhanced structural contacts between the N-terminus (≈ residues 1-8) and those near the CHC (Table S3). Specifically, residues ≈ 12-21 in cluster 5, and ≈ 10-20 in cluster 8 engage in interactions with the N-terminus. The contact maps of clusters 4 and 6 (Figure S8) also indicate enhanced ordering near the N-terminus of A*β*40. However, these features are less prominent than those in clusters 5 and 8, and the contacts tend to be more short-ranged (between residues *i* and *i* + 3). Cluster 12 consists of a nearly equal population of RC-like conformations, and those that have a number of long-range contacts between the N-terminus and residues ≈ 15 to 35 (Figure S8 and Table S3). Both simulations (22, 23, 68) and experiments (62, 63, 69, 70) have previously underscored the importance of residual structure in this region (particularly, LVFFA, residues 17 to 21), and the potential role in dictating aggregation kinetics. Overall, clusters containing exclusively RC-like states account for 32% of the equilibrium population. The cumulative population of clusters having a mixture of RC-like conformations, and those having residual structuring near the N-terminus is 47.3%. Taken together, ≈ 80% of the conformational ensemble is predominantly RC-like.

### Fingerprints of the fibril structure

Despite the dominance of RC-like conformations in the equilibrium ensemble, the essential fingerprints of the fibril state can be found in a few conformational clusters. Importantly, at the distance cutoff used for structural clustering, none of the constituent structures in these clusters resemble random-coils. Previous studies have unequivocally shown that the D23-K28 salt-bridge, which is a key structural element of mature A*β*40 fibrils, modulate the early events of amyloid aggregation (20, 21, 31). We find clear-cut signatures of these contacts in the structures in cluster 9 (Figure 5(b)), which is associated with an equilibrium population of 7.7%. In addition to the presence of E22/D23-K28 salt bridges, a majority of structures also have hydrophobic contacts between V24 and N27, which stabilize the VGSN turn. However, the N-terminus and C-terminus segments remain mobile, and hence do not correspond to hairpin-like conformations. In contrast, as evidenced by their relatively low *R*_*g*_ and *R*_*ee*_ (Figure S7) and prominent off-diagonal elements in the contact map, structures within cluster 11 (population 10%) have a high tendency to form hairpin-like structures via association of the two ends (Figure 5(d)). Many of the conformations within cluster 10 (population 3.2%) are ordered, and exhibit signatures of the fibril state (Figure 5(c)). In these structures, contacts are formed between the CHC and residues 25-40.

### A*β*42

Conformations belonging to clusters 2,3, 8 and 9 consist of exclusively RC-like structures. Some clusters identified for A*β*42 consist of an admixture of RC-like conformations, and ones with different extent of local structuring (Table S3). In cluster 1, nearly two-thirds of the constituent structures feature contacts between he central region (≈ residues 22 to and the C-terminus (Figure S9). Structures within cluster 10 are mostly semi-compact, and around 20% of these exhibit some tendency of structuring near the N-terminus by forming contacts with residues ≈ 20-30 (Figure S9 and Table S3). Conformations within cluster 11 are mostly disordered, and are typically RC-like, although a few of them exhibit structuring near the central region (Figure S9). Together, clusters having predominantly RC-like features (albeit with some degree of local structuring) contribute around 62.1% to the equilibrium population (Figure 5(a)). In addition to those having RC-like features, there are other clusters in the conformational ensemble of A*β*42, which exhibit structural features of different experimentally characterized fibril morphologies. We describe them below.

### Fingerprints of the S-bend fibril structure

Recently, NMR (44, 46) and cryo-EM (71) experiments have identified the “S-bend” motif as a building block of the A*β*42 fibril structure. The S-bend structure, which apparently is not found in A*β*40, results from enhanced ordering near the Cterminus of A*β*42, and is characterized by a close contact between residues K28 and A42. Although not immediately discernible from the contact maps, many of the conformations within clusters 4 and 5 exhibit the characteristic S-bend motif, stabilized by a close contact between residues K28 and A42 (Figure 5). These clusters exhibit enhanced ordering near the C-terminus, but differ somewhat in the specific residues involved in contact formation. In cluster 4 (Figure 5(e)), the contacts are formed between the CHC and residues 18-21, while in cluster 5 (Figure 5(f)) residues 15-21 form contacts with the CHC. Conformations similar to these have been previously identified by Garcia and coworkers using all-atom simulations in conjunction with spectral clustering algorithms (59).

### Fingerprints of the U-bend fibril structure

Besides the S-bend topology, A*β*42 also forms fibril structures in which the peptide adopts the canonical U-bend (72) (similar to the repeating unit of A*β*40 fibrils). Clusters 6 (Figure 5(g)) and 7 (Figure 5(h)) exhibit typical *β*-hairpin like structures, and consist of the E22/D23-K28 salt bridges, as well as the VGSN turn in a large fraction of the constituent structures. These clusters contribute 9.7% and 10.2% to the equilibrium population. In cluster 6, the hairpins are formed via the association between residues ≈ 25-35 and the N-terminus, whereas hairpins within cluster 7 involve contacts with the CHC.

From our clustering analyses we draw important insights regarding the sequence-specific heterogeneities of A*β*40 and A*β*42 monomers: (a) For both the sequences, clusters that are predominantly RC-like, or have RC-like structures in an admixture with conformations having some degree of residual structuring dominate the equilibrium population. Notably, these clusters do not exhibit any of the characteristic structural features found in the fibril. (b) Interestingly, despite having identical sequence for the first forty residues, the structural features near the termini are distinct for the two A*β* monomers. A*β*40 has a structured N-terminus, while A*β*42 exhibits enhanced structuring near the C-terminus, which is consistent with previous studies (19, 42, 48, 73, 74). As argued in a previous work (42), in A*β* monomers, there is a tug of war between the C-terminus and the N-terminus for making contacts with the CHC. In case of A*β*40, the N-terminus wins, because unlike A*β*42, it does not contain an extra hydrophobic patch formed by I41 and A42 to induce structuring near the C-terminus. Structuring near the N-terminus of A*β*40 could also have implications in the subsequent assembly process, and in determining the fibril morphology. As shown in a recent study (75), a cryo-EM A*β*40 fibril structure derived from a meningeal Alzheimer’s tissue is characterized by pronounced arches at the two ends (most notably at the N-terminus). (c) Although masked in ensemble-averages, clustering analyses reveal that the aggregation-prone structures (N* states) are present in the spectrum of conformations populating the ensemble, in agreement with previous studies (21, 32, 34, 36). For A*β*40, we find evidence of formation of E22/D23-K28 saltbridges, as well as antiparallel *β*-hairpins, and within the A*β*42 ensemble, we find signatures of both the classical *β* hairpins characterized by U-bends, and the recently discovered S-bend motifs.

### Linking the population of N* states to aggregation propensity

The analyses based on hierarchical clustering show that fibril-like or aggregation-prone conformations (*N*\*) are populated in the monomer ensemble of the A*β* peptides. However, in order to quantify their population, *p*_*N*\*_, a stringent order parameter, comparing the putative N* conformations identified using the cluster analysis with the fibril structure, is required. In other words, only a subset of conformations belonging to the N* basin of attraction has the propensity to aggregate. We estimate *p*_*N*\*_ based on a geometric criterion. Because the conformations belonging to the *N*\* state should be similar to the structure in the fibril, we use the fibril structures as reference states. We used the structural overlap parameter, *χ*_*fib*_(*t*), between a monomer conformation generated in the simulations at time *t*, and the monomer in the experimental fibril structure to characterize the *N*\* states for both the peptides (see Methods for further details). For A*β*40, we used the striated fibril structure determined by Tycko and coworkers (76) as the reference. ForA*β*42, we used the two structures reported by Riek and coworkers: the U-bend topology, which is very much similar to the repeating unit found in A*β*40 fibrils (72), and the S-bend motif (44).

The distributions of *χ*_*fib*_ are shown in the top panel in Figure 6. The distributions for A*β*40 and A*β*42 nearly overlap when the U-bend (or striated) structure is chosen as a reference. This suggests that the population of fibril-prone structures exhibiting the canonical *β*-hairpin motif between opposing strands, is approximately similar between the two sequences. In contrast, the conformational ensemble of A*β*42 seems to bear a higher overall resemblance to the S-bend motif.

The *N*\* state is hidden in the tail of these distributions (Figure 6(a)). To identify the approximate location of the basin consisting of the aggregation prone conformations, we carried out restrained simulations (see the SI for details) starting from the monomer structure present in the experimental fibril morphologies of A*β*40 and A*β*42. The 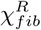 distributions are calculated from the restrained simulations reflect the conformations within the *N*\* basin of attractions that could readily aggregate (purple curves in Figures 6(b), 6(c) and 6(d)). We find that the basin of aggregation-prone structures is approximately centered at *χ*_*c*_ = 0.28 for both A*β*40 and A*β*42. Hence, for both the sequences, we collectively define *N*\* states as the ensemble of conformations with *χ*_*fib*_ ≥ *χ*_*c*_. Therefore, *p*_*N*\*_ we calculated using,

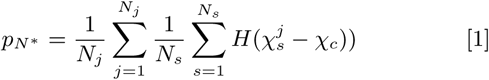

where 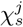 is the structural overlap for the *s*^*th*^ snapshot in the *j*^*th*^ trajectory; *N*_*s*_ denotes the number of snapshots in each trajectory; and *N*_*j*_ denotes the number of independent trajectories; and *H* is the Heaviside function. Using Eq. 1, we estimated that *p*_*N*\*_ is 0.45% for A*β*40. For A*β*42, *p*_*N*\*_ is estimated to be 1.45% when the U-bend topology is used as a reference, and 2.17% when the S-bend motif is used as a reference. These numbers fall within the same range as previous estimates of *p*_*N*\*_ at physiological conditions for the src SH3 (33) domain, which is a globular protein.

As indicated by the presence of a minor peak near *χ*_*c*_, only a subpopulation of conformations within the clusters (Figure S10), which exhibit structural signatures of the fibril state, meet the strict geometric definition (Eq. 1) for the conformations within the N* state that are likely to aggregate. This observation implies that although contact maps corresponding to various sub-ensembles could be useful in discerning the presence of aggregation-prone species, they are probably too coarse as metrics for quantitatively measuring *p*_*N*\*_.

In our previous work (32), we proposed that the time scale of fibril formation, which should be thought of an approximate estimate of the mean first passage time for the monomer to reach the structure in the fibril (U-bend or S-Bend), is related to *p*_*N*\*_ as,

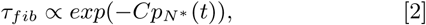

where the population of the *N*\* state is expressed as a percentage. The expression in Eq.2 does not account for the multiplicity of events that are involved in fibril formation. Using *C*=1 for both the peptides, and the estimated values of *p*_*N*\*_, we predict that *τ*_*fib*_ for A*β*42 is approximately 24 times smaller than A*β*40. Our prediction is in accord with recent experiments (39), which show that the fibril formation rate of A*β*42 is about one order of magnitude greater than A*β*40. Interestingly, the one order of magnitude difference cannot be rationalized if only the U-bend or striated conformations are identified with the *N*\* state. In this case, the aggregation of A*β*42 is only twice as fast. This implies that the enhanced ordering near the C-terminus, which stabilizes the S-bend motif, provides an alternate route for fibril formation in A*β*42, and could possibly explain why it is more aggregation-prone despite being present in a much lower concentration than A*β*40 in the plasma of cells.

## Discussion

We used the SOP-IDP model to characterize the conformational ensembles of A*β*40 and A*β*42 monomers in order to quantify the role their excited states play in controlling fibril formation. The ensemble-averages of many properties are practically identical for the two sequences. They exhibit identical Flory scaling behavior for the dependence of the inter-residue distance as function of sequence separation, indicating that the ground state of both the peptides are best described as random coils. The chemical shifts for the peptides are also similar. Clustering analyses revealed the details of sequence-specific heterogeneity. As evidenced by the inter-residue contact maps for individual clusters, both the peptides access an array of different structures at room temperature. Specifically, the A*β*40 monomer has a measurable probability of adopting a structured N-terminus, while the C-terminus of A*β*42 is ordered to some extent. For both the sequences, we also find prominent fingerprints of the existence of aggregation-prone (excited) conformations (N* states), which are only sparsely populated. The N* states share a number of features in common with the monomer structures in the fibrils (Figure 7). By using the differences in the population, *p*_*N*\*_, of the N* states (or equivalently the free energy difference, Δ*G*_*N*\*_ with respect to the disordered ground state) between the two peptides in an empirical relation connecting *p*_*N*\*_ to fibril elongation rates (Eq. 2), we rationalize the higher value of *τ*_*fib*_ in A*β*40 relative to A*β*42.

**Fig. 7.**
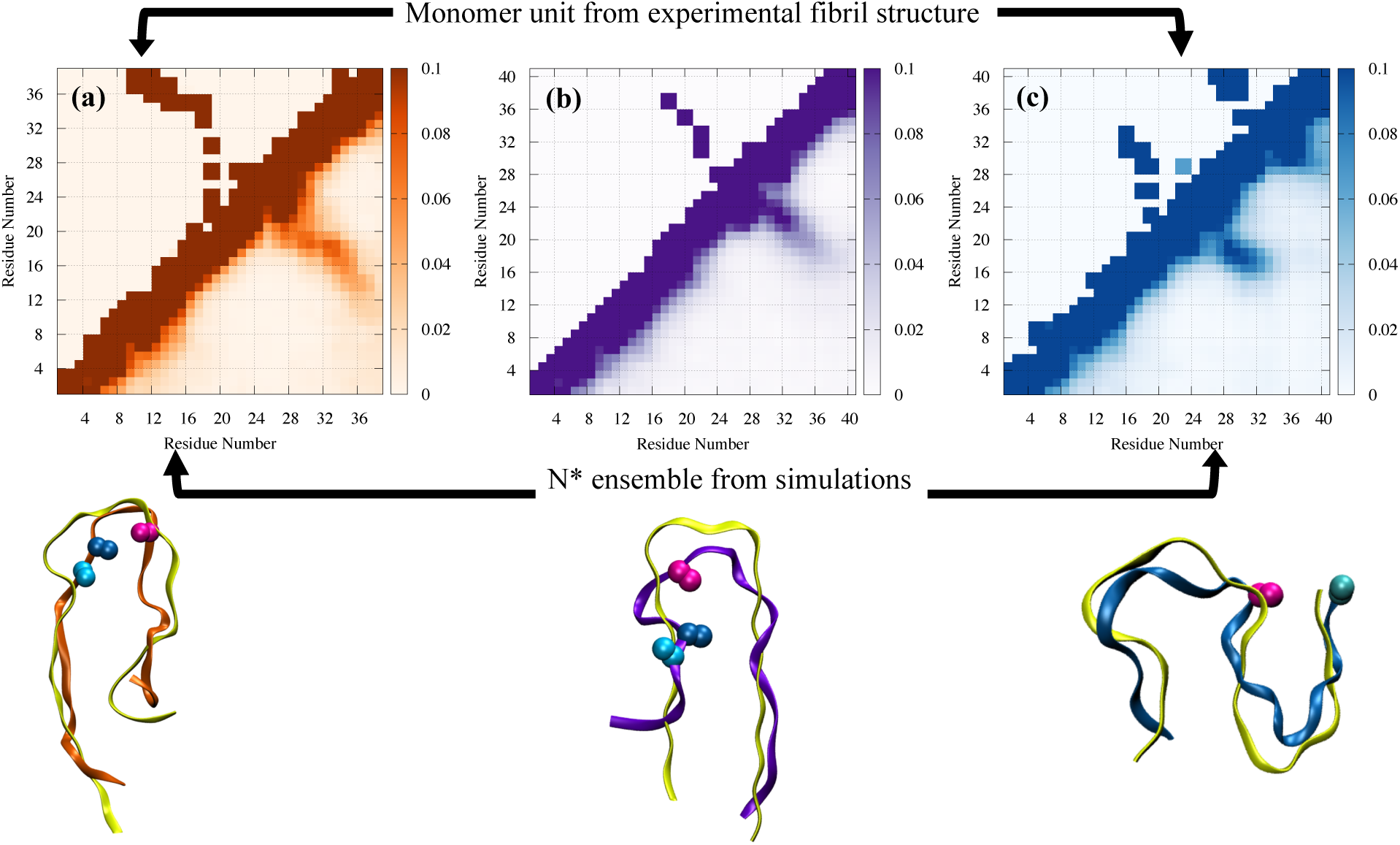
Contact maps for the *N*\* ensemble identified form simulations (lower triangle), and the monomer unit of the experimental fibril structure (upper triangle). (a) *N*\* ensemble of A*β*40, consisting of hairpin-like configurations, stabilized by the VGSN turn, and D23-K28 salt bridge. As is evident, these structural features are also found in the A*β*40 fibril structure. At least two structurally distinct N* states exist for A*β*42, as shown in (b) and (c). (b) Hairpin-like structure consisting of a U-bend near the VGSN region, and stabilized by the D23-K28 salt bridge. (c) S-bend motif stabilized by a contact between K28 and A42. Below the contact maps, a representative structure from the *N*\* ensemble is shown superimposed on the monomer unit from the fibril structure (shown in yellow). For clarity, we only show residues that are part of the fibril core (10-40 for A*β*40, and 17-42 for A*β*42) are shown.

The propensity to form *β* structures in the spectrum of monomers could be used as a marker for protein aggregation. Indeed, the slightly upshifted values of the chemical shifts for some residues relative to the ensemble-averages (Figure S11) does imply that the overall *β* propensity is somewhat enhanced in the *N*\* ensemble. Nonetheless, the *N*\* state retains some degree of disorder, which is essential for the rapid conformational switching between the different structures. It is likely the complete structural ordering into *β*-sheets does not occur until the later stages in the aggregation cascade (formation of critical nuclei or protofibrils), when “crowding” effects due the presence of neighboring monomers resulting in favorable inter peptide interactions compensate for the loss of monomer conformational entropy.

### Link between N* states and sequence and environment dependent aggregation propensities

The idea that propensities of amyloid assembly could be discerned from the population of aggregation-prone high free energy monomer states, *p*_*N*\*_ has important implications. (1) Because the *N*\* state is separated from the ground state by a free energy gap, Δ*G*_*N*\*_, it follows that *p*_*N*\*_ is vanishingly small if Δ*G*_*N*\*_ */k*_*B*_ *T* ≫ 1. In this scenario, aggregation is unlikely. (2) The aggregation propensity, which is dictated by *p*_*N*\*_, is slave to external conditions since Δ*G*_*N*\*_ can be modulated by varying the temperature, pH, or crowder concentration (14). This implies that the full sequence-dependent folding landscape of the monomer has to be determined in order to estimate *p*_*N*\*_. A corollary of our finding is that it is difficult, if not impossible, to predict aggregation propensities from sequence gazing alone (77). (3) Many proteins can be made to aggregate by arranging conditions such that the Δ*G*_*N*\*_ */k*_*B*_ *T* ≫ 1 is not too large, which tidily explains why even the helical protein myoglobin can form fibrils under certain conditions (78). Sequences associated with higher values of *p*_*N*\*_ for a given set of external conditions, are therefore likely to have higher rates of aggregation. For instance, it is known that introduction of a lactam bridge near the VGSN turn region of A*β* enhances the kinetics of fibril formation in A*β*40 by a factor of 1000 (79), which was explained by substantial increase in the population of aggregation-prone *N*\* states (21).

### Generality of the *N*\* theory

Even if the lowest free energy state is not a random coil but is a globular protein, aggregation to a fibril is possible only if the *N*\* state is populated to some extent. This was illustrated by analyzing the experiments on the aggregation of the SH3 domain, a small globular protein. By using relaxation dispersion NMR experiments (80) it was shown that under ambient conditions a very low percentage of excited states is populated. Using simulations we showed that *p*_*N*\*_ ≈ 2%, which allowed us to estimate the fibril elongation rate nearly quantitatively (33) using Eq.2. Significantly, the *N*\* determined in simulations was structurally identical to the one determined using NMR (see Fig. S7 in (33) in which the two structures are super imposed). The earlier study (33) affirmed that even for globular proteins *p*_*N*\*_ provides a good estimate of fibril formation times and that the aggregation prone structures have similarities to the monomer in the fibril. Thus, it appears that the empirical theory based on the *N*\* concept might be general. Needless to say, additional studies are needed to further validate the the theory that excited states of amyloid forming proteins could provide estimates of relative fibril formation times.

**Fig. 8.**
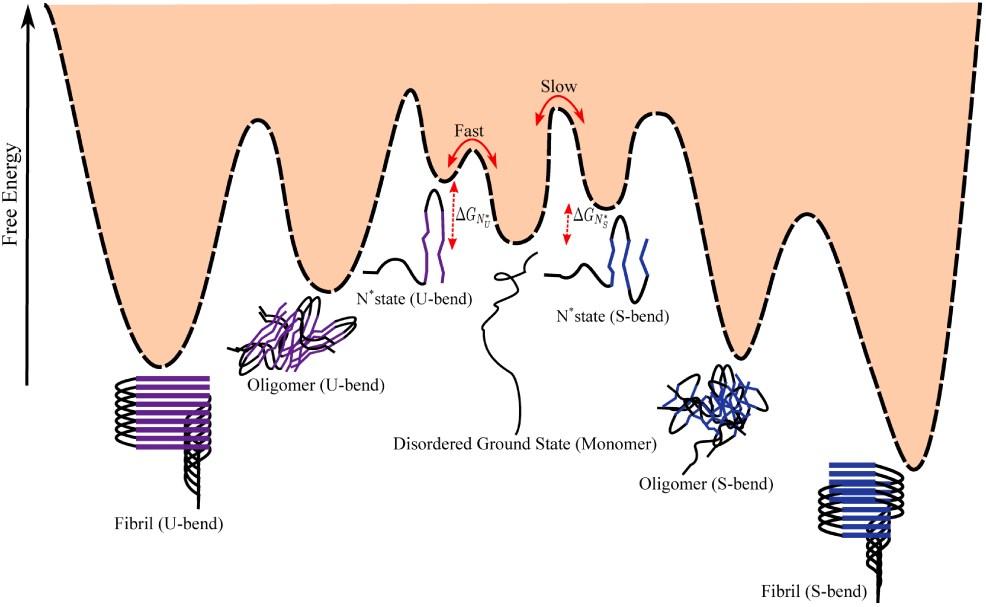
A schematic free energy landscape illustrating the aggregation cascade that lead to the formation of fibrils, starting with the transition from the disordered ground state to the *N*\* ensemble at the monomer level, and ultimately the formation of fibril structures from an oligomeric assembly of *N*\* states via different growth mechanisms. For A*β*42, the seeds of polymorphism are encoded in the structurally diverse *N*\* states. Furthermore, the aggregation cascade seems to be in harmony with the Ostwald’s rule. For A*β*42, this would imply that 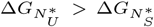, and hence transition to the U-bend conformation is likely to be faster.

### Polymorphism and Ostwald’s rule

The multiplicity of *N*\* states (Figure 7), as revealed in the heterogeneity of their conformations, implies that there could be several polymorphic fibril structures. The formation of such structures depends on the conditions of fibril growth. An implication of our study is that existence of polymorphism, including the potential role of water (81), are also encoded in the excitation spectra of the monomer.

We illustrate some of the concepts underlying fibril polymorphism through a schematic free energy landscape (Figure 8). It is likely, especially in A*β*42, which has more than one fibril structure, that the appearance of distinct polymorphs follows Ostwald’s rule. Indeed, a recent study demonstrated how Ostwald’s rule, well-known in the formation of crystal polymorphs, is manifested during the supramolecular assembly of synthetic polymers (82). Ostwald’s rule affirms that the least stable polymorph would form first, followed by a subsequent rollover to the more stable form. We predict that for A*β*42, transition from the disordered ground state to the U-bend topology is likely to be faster, with the S-bend topology only appearing on longer observation time scales. This prediction, which follows from the relative free energy difference between the two excited states of the A*β*42 monomer (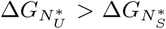, with 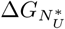 and 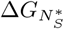 denoting the free energy difference of the U-bend and the S-bend motifs with respect to the disordered ground state), could be tested using kinetic simulations. It follows (see Figure 8) that formation of various polymorphic structures could be under kinetic control, which was shown in a prescient study sometime ago (83).

We also speculate that the signature of fibril polymorphism formed under a specific set of external conditions is determined fairly early during the aggregation cascade, perhaps even at the level of monomer fluctuations. The competition between different *N*\* ensembles could modulate the equilibrium population of the distinct polymorphs. It also seems highly unlikely that different fibril polymorphs could ‘directly’ interconvert from one form to another, without undergoing step-wise disassembly into oligomers and monomers. Thus, once formed the various polymorphic structures are trapped implying that formation of distinct fibril structures are under kinetic control (83). The landscape underlying aggregation can of course be tuned by varying the external conditions, to shift the balance between the *N*\* states, which ultimately regulates the polymorphic fate of amyloid fibrils.

### Materials and Methods

The A*β*40 and A*β*42 peptides are modeled using a modification of the recently introduced Self-Organized Polymer (SOP) IDP model (abbreviated as SOP-IDP), which quantitatively and in unprecedented detail reproduces the scattering profiles of a diverse range of IDP sequences of varying lengths, sequence composition, and charge densities (52). We calculated the thermodynamic quantities using trajectories generated in low friction Langevin dynamics simulations, which enhances conformational sampling (84). The simulations were used to calculate a number of observables for the A*β*40 and A*β*42 peptides, which were directly compared to experiments in order to validate the SOP-IDP model. Details of the simulations and analyses of the simulations may be found in the Supplementary Information (SI).

## ACKNOWLEDGMENTS

We are grateful to Upayan Baul for assistance in the initial stages of this work. The comments of Robert Tycko and the encouragement of Tuomas Knowles are greatly appreciated. We also acknowledge Mauro Mugnai and Abhinav Kumar for several fruitful discussions. The authors acknowledge the Texas Advanced Computing Center (TACC) for providing the necessary computing resources. This work was supported by a grant from the National Institutes of Health (GM - 107703).

## Supporting Information Text

### Computational Methodology

#### The SOP-IDP model

The A*β*40 and A*β*42 peptides are modeled using a modification of the recently introduced Self-Organized Polymer (SOP) IDP model (abbreviated as SOP-IDP), which quantitatively and in unprecedented detail reproduces the scattering profiles of a diverse range of IDP sequences of varying lengths, sequence composition, and charge densities (1). In the SOP-IDP model, each amino-acid residue is represented using two interacting sites: a backbone bead (BB) centered on the *C*_*α*_ atom, and the other is a side-chain bead (SC) centered on the center-of-mass of the side-chain (Figure S1). The SOP-IDP coarse-grained energy function is:

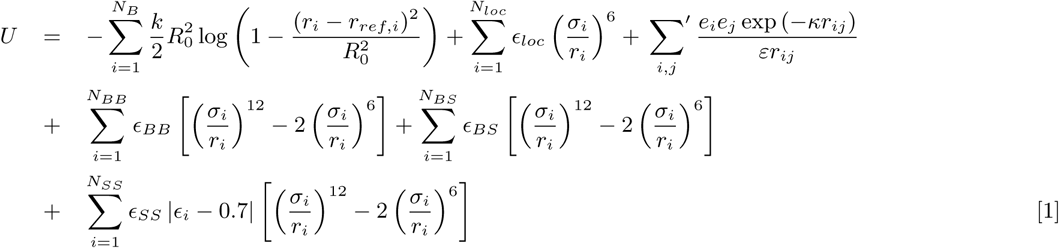

The first term in Eq. 1, with *r*_*ref,i*_ being the equilibrium distance between bonded moieties (two consecutive backbone beads and side chains that are connected to backbone beads), is the finitely extensible nonlinear elastic (FENE) potential, which accounts for the chain connectivity. The second term denotes the repulsive excluded volume interactions, which prevents unphysical overlap between beads. The third term describes the electrostatic interactions between all the pairs of charged residues expressed in terms of the standard Debye-Hückel relation. The final three terms in the energy function account for the sequence-specificity, and describe the backbone-backbone (BB), backbone-sidechain (BS), and sidechain-sidechain (SS) interactions, respectively. The parameter *ϵ*_*i*_ in Eq. 1 is based on the Betancourt-Thirumalai statistical potential (2). There are only three crucial parameters in the SOP-IDP model. They are *ϵ*_*BB*_, *ϵ*_*BS*_, and *ϵ*_*SS*_. Their values were determined using a learning procedure described elsewhere (1). The values of the different parameters used in the SOP-IDP model are given in Tables S1 and S2.

**Fig. S1.**
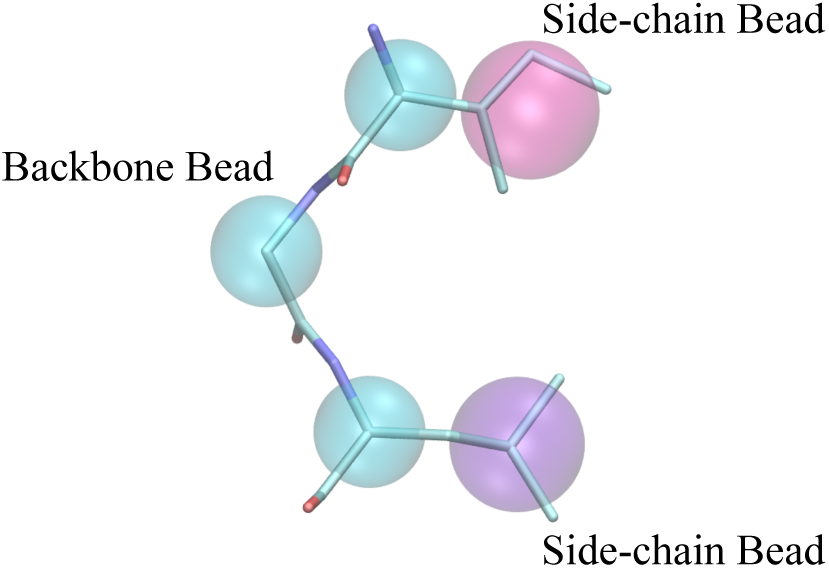
Schematic illustration of the coarse-grained representation of polypeptide chains in the SOP-IDP model. The backbone bead (shown in cyan) is centered on the *C*_*α*_ atom. The side-chain beads (shown in magenta and purple colors illustrate the sequence-specific differences) are centered on the center-of-mass of the side-chains.

#### Simulations

We calculated the thermodynamic quantities using trajectories generated by Langevin dynamics simulations, where the motion of each bead *i*, is described by:

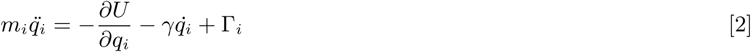

where *m*_*i*_ is the mass of the bead, *q*_*i*_ denotes the coordinate, *γ* is the friction coefficient, and Γ_*i*_ is a Gaussian random force, which satisfies ⟨Γ_*i*_(*t*)Γ_*j*_ (*t*′)⟩ = 6k_*B*_ T*γδ*_*ij*_ *δ*(*t* − *t*′). To enhance the conformational sampling (3), the simulations were performed in the low friction regime, corresponding to a solvent viscosity of 10^−5^ Pa.s (which is around 1% of the viscosity of water). In our model, the average mass, *m*, of a bead is 53 g/mol, while the typical energy scale, *E*, and length scale *a*, are 1 kcal/mol, and 1 Å, respectively. From these, the natural unit of time in the low friction regime (Eq. 2) is estimated to be 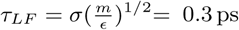. The equations of motion were integrated using a modified version of the velocity-Verlet algorithm (3), employing a time-step of 0.05*τ*_*LF*_. To obtain meaningful statistics for thermodynamic observables, simulations were carried out from ten different initial conditions, for at least 2× 10^9^ steps. All the simulations were performed at 298 K.

**Table S1.** Energy function parameters for the SOP-IDP model.

| Parameter | Value |
| --- | --- |
| $R_0$ | 2.0 Å |
| $k$ | 20.0 kcal/(mol Å<sup>2</sup>) |
| $\epsilon_{loc}$ | 1.0 kcal/mol |
| $\epsilon_{BB}$ | 0.12 kcal/mol |
| $\epsilon_{BS}$ | 0.24 kcal/mol |
| $\epsilon_{SS}$ | 0.18 kcal/mol |

**Table S2.** Parameters for the coarse-grained beads used in the SOP-IDP model.

| Residue | Radius (Å) | charge (e) |
| --- | --- | --- |
| Gly | 0.0 | 0.0 |
| Ala | 2.52 | 0.0 |
| Val | 2.93 | 0.0 |
| Leu | 3.09 | 0.0 |
| Ile | 3.09 | 0.0 |
| Met | 3.09 | 0.0 |
| Phe | 3.18 | 0.0 |
| Pro | 2.78 | 0.0 |
| Ser | 2.59 | 0.0 |
| Thr | 2.81 | 0.0 |
| Asn | 2.84 | 0.0 |
| Gln | 3.01 | 0.0 |
| Tyr | 3.23 | 0.0 |
| Trp | 3.39 | 0.0 |
| Asp | 2.79 | -1.0 |
| Glu | 2.96 | -1.0 |
| His | 3.04 | 0.0 |
| Lys | 3.18 | 1.0 |
| Arg | 3.28 | 1.0 |
| Cys | 2.74 | 0.0 |
| backbone | 1.90 | 0.0 |

#### Restrained simulations - identification of aggregation-prone N* conformations

In order to determine the stringent criterion of aggregation-prone conformations we performed restrained simulations starting from the monomers in the A*β*40 and A*β*42 fibril structures. Each restrained simulation was carried out for 10^8^ time steps. For A*β*40, we used the fibril structure reported by Tycko and coworkers (4) (PDB ID: 2M4J) as the initial conformation in the restrained simulations. To maintain the overall fibril-like topology throughout the simulations, a harmonic restraint was added between the C_*α*_ atoms of residues 13 and 38. For A*β*42, the initial structures for the restrained simulations were based on two different fibril morphologies identified using solution-NMR, the U-bend structure (5) (PDB ID: 2BEG) and the S-bend structure (6) (PDB ID: 2NAO). For the U-bend structure, the distance between the C_*α*_ atoms of residues 17 and 41 were constrained using a harmonic potential. To maintain the S-bend topology during the simulations, two harmonic distance restraints were necessary: between the C_*α*_ atoms of residues 17 and 33, and residues 28 and 41. In all these simulations, a strength 50 kcal/mol Å^−2^ was found to be an optimal value for the distance restraints, as it maintained the overall topology of the initial structure, without introducing spurious distortions. The distribution of 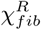 serves as a reference in determining the aggregation-prone N* conformations.

### Analyses

#### End-end distance (*R*_*ee*_) distributions for standard polymer models

The theoretical predictions for the *R*_*ee*_ distributions of a self-avoiding random walk (SAW) and Gaussian chain are given by: 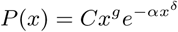 for number of residues, *N*_*tot*_ ≫ 1 (7). Here, 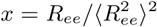, and *C* is a normalization constant. The exponent *δ* = 1/(1 − *ν*) (with *ν* ≈ 0.588 for a SAW and 0.5 for a Gaussian chain) accounts for the decayJ of *P* (*x*) for *x* > 1. The Jcorrelation hole exponent, *g* ≈ 0.28 for a SAW, and *g* = 0 for a Gaussian chain. The conditions 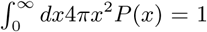 and 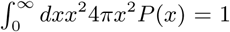 allow the determination of *C* and *α*. For a Gaussian chain *C* = (3/2*π*)^3^/2 and *α* = 1.5, whereas for a SAW *C* ≈ 0 278 and *α* ≈ 1 206. In the limit of large *N*_*tot*_, the interaction between side-chains does not affect the shape of *P*(*x*) and, hence *P*(*R*_*ee*_) is expected to display the universal behavior. Deviations from the universal theoretical predictions likely occurs due to finite size of the polypeptide chains and/or is as consequence of short-range intra-peptide interactions. For the A*β* peptides considered here, finite-size effects are substantial, and hence significant deviation from the theoretical curves is observed (Figure 1(c) in the main text).

#### Contact Maps

Two residues *i* and *j* are assumed to be in contact if the distance between their corresponding side-chains are less than or equal to 0.8 nm. The time-dependent inter-residue matrices are converted to probability contact maps *p*_*ij*_ (*t*) using the logistic function,

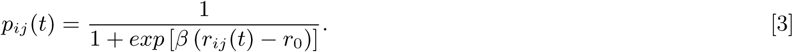

In Eq. 5, *β* = 500 nm^−1^, and *r*_0_ = 0.8 nm. The ensemble-averaged probability maps, 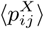 (where X= A*β*40 or A*β*42) are computed by taking an average over all the snapshots *N*_*s*_ in trajectories *N*_*j*_.

#### Secondary Structure Analysis

The secondary structures of each amino acid residue for a given coarse-grained (CG) conformation were assigned using the PCASSO program developed by Brooks and coworkers (8). In this scheme, first a training set consisting of proteins having a wide range of secondary structures is used to optimize a random forest classifier using machine learning, and subsequently secondary structure propensities for a protein sequence are predicted based on the positions of the C_*α*_ atoms. Specifically, the secondary structure assignment in PCASSO uses a three-class system, which identifies residues as a helix (h), *β*-sheet (s), or turn/coil configuration (c) (8). No distinction is made between turn and coil states. As noted by the authors (8), the predictions of PCASSO are in excellent agreement with other schemes, such as DSSP and STRIDE, which, however, require hydrogen-bonding information, in addition to coordinates of all the atoms.

#### Calculation of Chemical Shifts

The chemical shifts were calculated from the CG trajectories using the LARMOR-C_*α*_ formalism (9). Briefly, this method uses a number of geometrical features based on C_*α*_-C_*α*_ distances, and is trained using a random forest classifier on the RefDB database, consisting of proteins for which both high resolution X-ray structures, and NMR chemical shifts are available. Here, we report the chemical shift for the C_*α*_ in residue *i* in terms of secondary chemical shift (10), 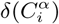, where:

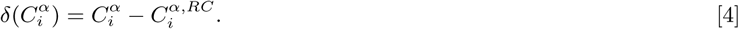

In Eq. 3, 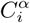 corresponds to the chemical shift computed using LARMOR-C_*α*_ and 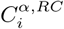 is the chemical shift for the reference random coil state in residue *i*. The 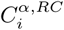 estimates are based on the values reported by Poulsen and co-workers (11, 12), and were obtained using the webserver maintained by the Bax group.

To assess the agreement between the calculated chemical shifts and experimental data, we computed the weighted root-mean-squared-error (*ζ*) given by:

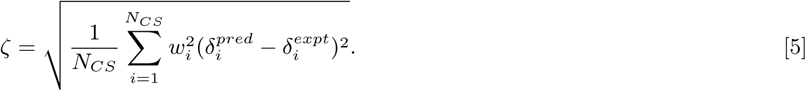

where, *w*_*i*_ is taken to be 0.965 based on the error associated with the estimation of *C*_*α*_ chemical shifts with the LARMOR-C*α*.

#### FRET efficiencies

We calculated the FRET efficiency from the time-series of rescaled end-end distance, 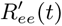 using the equation:

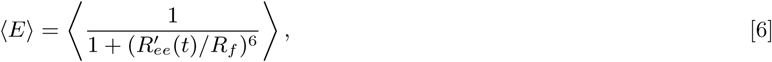

where, *R*_*f*_, the Forster radius is taken to be 5.2 nm. The angular brackets denote the average over the distributionof end-to-end distances obtained from simulations. Following previous studies,(13, 14) we account for the effect of the attached dyes in FRET experiments, by rescaling the end-end distance, *R*_*ee*_:

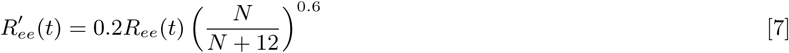

where *N* is either 40 or 42 depending on the peptide sequence.

#### Principal Component Analysis

In order to identify possible metastable states of the A*β*40 and A*β*42 peptides, we performed a principal component analysis (PCA) of the simulated trajectories in the space of the pairwise C_*α*_–C_*α*_ distances, *x*_*n*_. The covariance matrix, *C*_*ij*_ = ⟨(*x*_*i*_ − ⟨*x*_*i*_⟩)(*x*_*j*_ − ⟨*x*_*j*_⟩)⟩, is diagonalized to yield the eigenvalues, *λ*_*n*_, and eigenvectors, *V*_*n*_, which are the amplitudes and the directions of the collective motions.

#### Radius of Gyration, (*R*_*g*_)

The radius of gyration was calculated using:

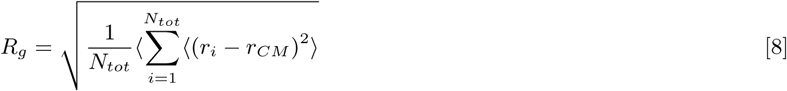

where *N*_*tot*_ is the total number of coarse-grained beads, and *r*_*i*_ are the coordinates of bead *i*, and *r*_*CM*_ is the center-of-mass coordinate, and ⟨…⟩ denotes the ensemble average.

#### Hydrodynamic Radius, (*R*_*h*_)

We calculated *R*_*h*_ using the standard polymer-physics equation:

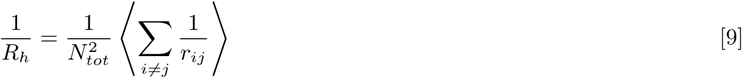

where *N*_*tot*_ is the total number of coarse-grained beads, and *r*_*ij*_ is the distance between beads *i* and *j*.

#### Hierarchical clustering of conformational ensembles

Our previous study (1) showed that IDP ensembles are highly heterogeneous, which is masked when only ensemble averages are calculated. The extent of heterogeneity of the different sequences are revealed using clustering techniques. To identify the representative conformations populating the A*β* ensembles at equilibrium, and decode the sequence-specific heterogeneity, we performed hierarchical clustering using the Ward’s variance minimization algorithm (15). The dimensionless distance metric, *D*_*ij*_, between two conformations *i* and *j* is denoted as:

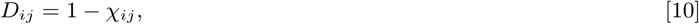

where *χ*_*ij*_, the structural overlap between conformations *i* and *j*, is given by:

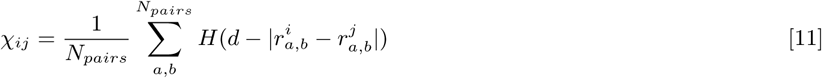

In Eq. 7, 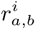 and 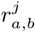 are the distances between coarse-grained sites *a* and *b* in snapshots *i* and *j*, respectively; *H* is the Heaviside step function, *d*=0.2 nm is the tolerance, which accounts for thermal fluctuations. The sum is taken over all the sites separated by more than two covalent bonds; *N*_*pairs*_ is the total number of such pairs. We used dendrograms to visualize the hierarchical organization of different conformations.

In order to unambiguously identify the long-range contacts in a given conformational cluster *Y*, we calculated the difference probability map 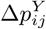, which is given by:

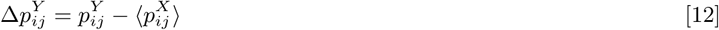

In Eq. 14, 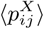 is the ensemble-averaged contact probability map for peptide *X* (where *X* = A*β*40 or A*β*42), and 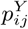 denotes the contact probability map for cluster *Y*. By definition, 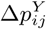 is bound between +1 and −1. Here, we restrict the contact probability maps to positive values of 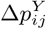 in order to discern how the propensity of a contact between residues *i* and *j* in a conformational cluster is enhanced with respect to the ensemble-average.

#### Identification of aggregation-prone (N*) states

Aggregation-prone conformations (the putative N* state) were identified from the conformational ensembles based on structural overlap with the monomer unit of the experimentally determined A*β*40 and A*β*42 fibril structures. We computed *χ*_*fib*_(*t*) using:

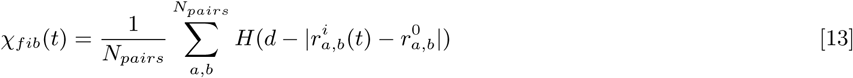

where 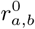 is the distance between sites *a* and *b* in the reference structure. Conformations which exhibit *χ*_*fib*_(*t*) ≥ *χ*_*c*_ are deemed to be aggregation-prone, and collectively defined as belonging to the N* state. For A*β*40, we chose the monomer unit from the solid-state striated fibril structure (PDB ID: 2M4J) reported by Tycko and coworkers (4) as the reference state. To identify aggregation-prone conformations in the A*β*42 ensemble, we used the monomer units from two topologically different fibril structures reported by Riek and coworkers using solution-NMR as references, the U-bend fibril structure (5), and a S-bend fibril structure (6).

**Fig. S2.**
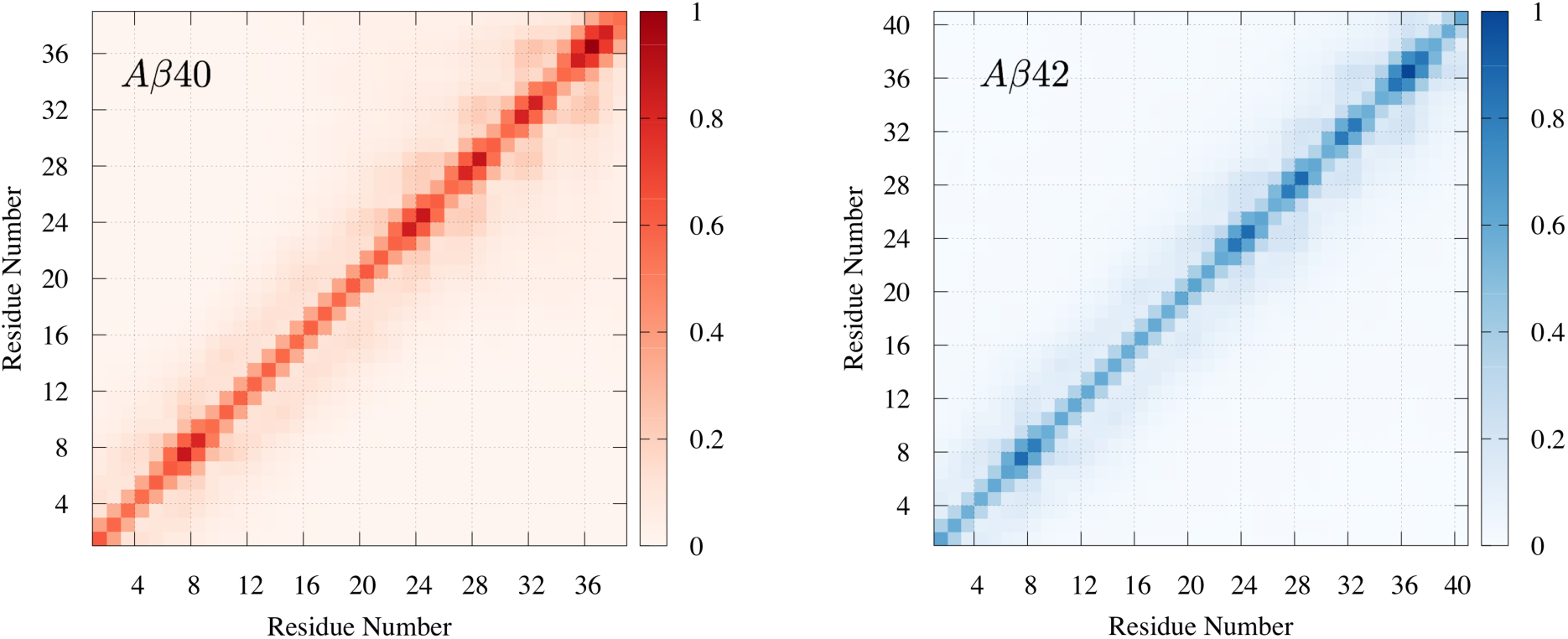
The ensemble-averaged contact maps for the A*β*40 and A*β*42 monomer ensembles. As is evident, the the monomers do not adopt long-lived secondary structures, and exhibit RC-like behavior.

**Fig. S3.**
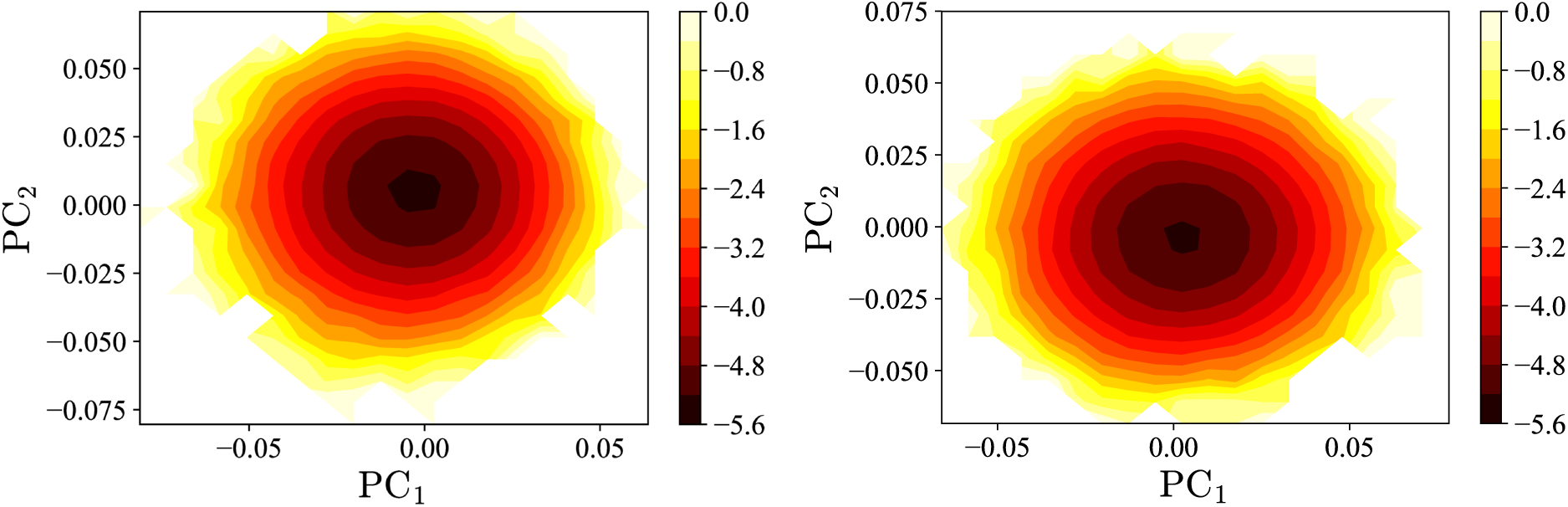
Free energy landscape of the A*β*40 (left) and A*β*42 (right) monomer projected onto the first two principal components, PC_1_ and PC_2_. The landscapes exhibit broad featureless contours, which exemplifies the disordered nature of the monomer ensembles. The color-bar denotes the free energy in kcal/mol.

**Fig. S4.**
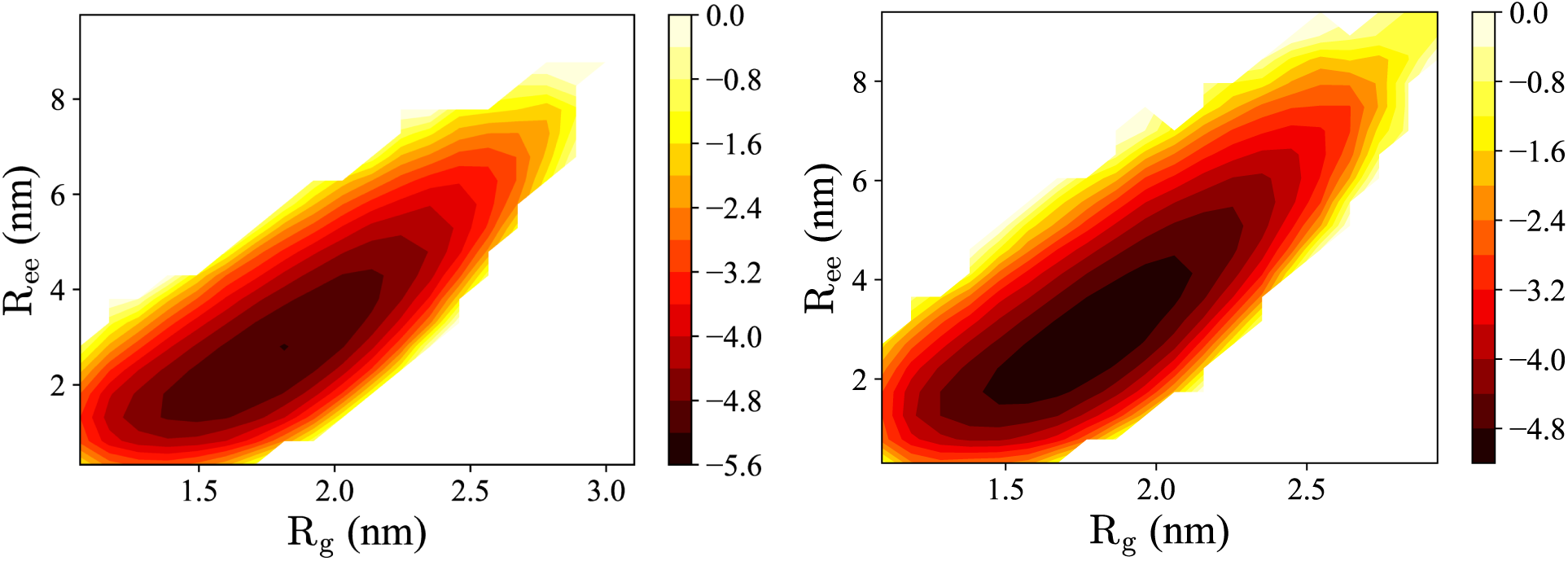
Free energy landscape of the A*β*40 (left) and A*β*42 (right) monomer projected onto the *R*_*g*_ and *R*_*ee*_ coordinates. Similar to Fig. S3, the landscapes are essentially featureless. The scales on the right have the same meaning as in Fig. S3.

**Fig. S5.**
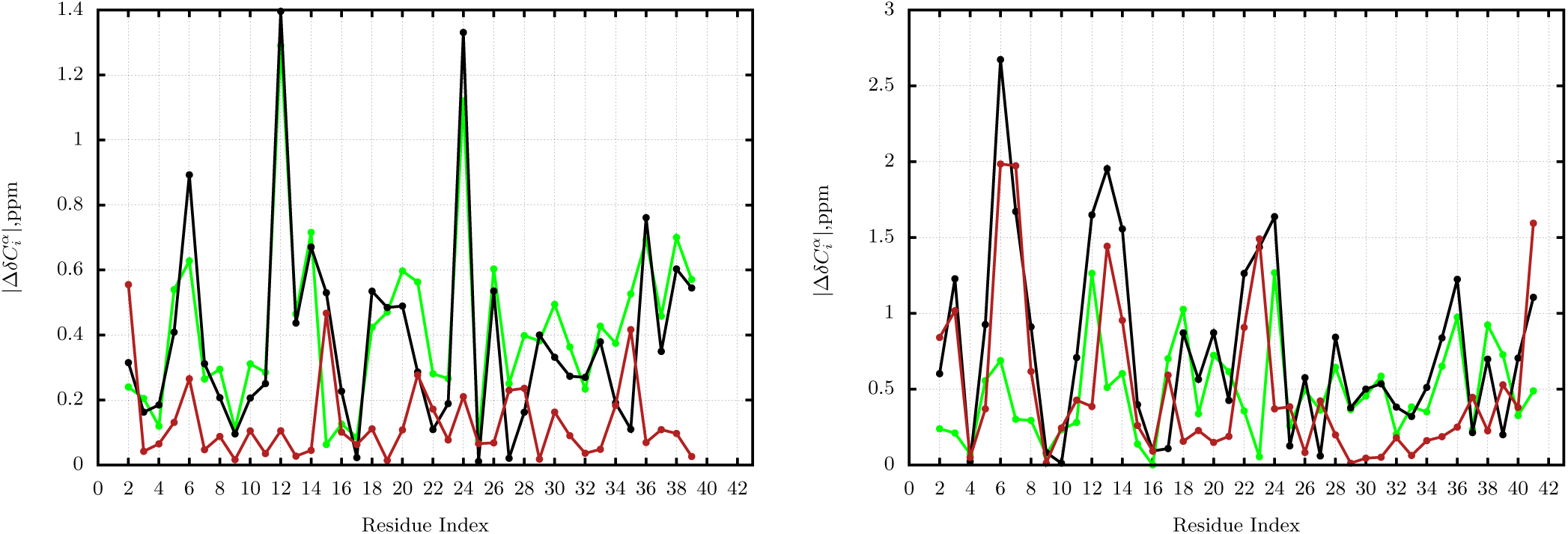
The green line denotes the residue-wise differences in the relative chemical shift, 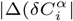 between simulations, and those measured by Bax and coworkers (16). The black line denotes 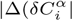 between the relative chemical shifts predicted by simulations and those measured by Zagorski and coworkers (17). The red line shows the differences between the two experimental measurements.

**Fig. S6.**
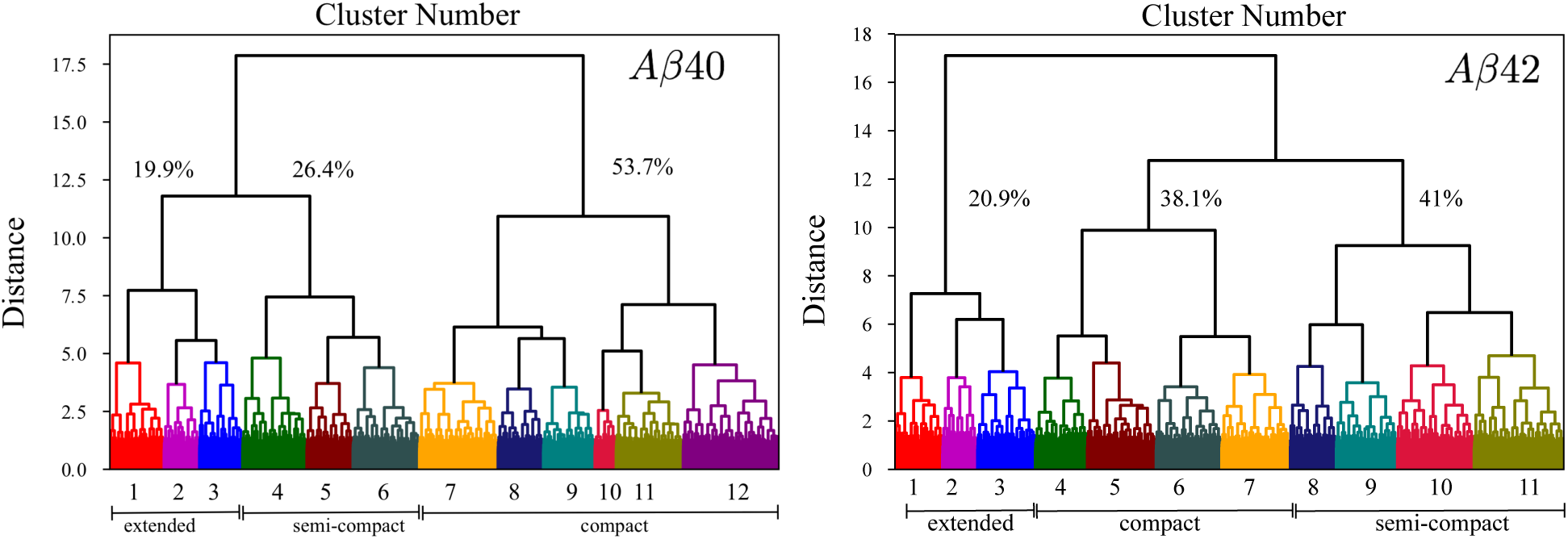
Hierarchical clustering of the conformational ensembles of A*β*40 and A*β*42. The hierarchical organization of the clusters is depicted in the form of dendrograms.

**Table S3.** The net populations of the different subclusters for A*β*40 and A*β*42 identified using hierarchical clustering. Some clusters contain a mixture of different structures, as described in the Table. *RC* denotes random-coil like structures, having featureless contact maps. *N*_*T*_ represents structures having residual ordering near the N-terminus. *C*_*T*_ correspond to structures having residual ordering near the C-terminus, and *M* denotes conformations having residual structuring near the middle of the sequence. Clusters with some of the structural signatures found in the fibril state are labeled *N*\*. Since RC-like features are present in all clusters except those characterized as *N*\*, we calculate the cumulative population of random coil states by summing over their populations. Overall, RC-like conformations contribute 80% to the equilibrium population of A*β*40, and 62% to the equilibrium population of A*β*42. The ≈ 20% of conformations of A*β*40 belong to the *N*\*basin of attraction. In the case of A*β*42, out of the 38% of conformations identifies as belonging to the *N*\* basin of attraction 20% (18%) of the conformations have U-bend (S-bend) type topology. From the conformations deemed to be in the *N*\* basin of attraction a strict criterion (see the main text) is used to determine the population of the aggregation-prone structures using Eq. (1) in the main text.

| Cluster Number | Aβ40 | Type | Aβ42 | Type |
| --- | --- | --- | --- | --- |
| 1 | 8.2% | RC (100%) | 7.2% | RC (27%) + $C_T$ (73%) |
| 2 | 5.3% | RC (100%) | 5.2% | RC (100%) |
| 3 | 6.5% | RC (100%) | 8.6% | RC (100%) |
| 4 | 9.6% | RC (70%) + $N_T$ (30%) | 7.7% | $N^*$ (100%) |
| 5 | 6.8% | RC (50%) + $N_T$ (50%) | 10.3% | $N^*$ (100%) |
| 6 | 9.9% | RC (60%) + $N_T$ (40%) | 9.7% | $N^*$ (100%) |
| 7 | 11.7% | RC (100%) | 10.2% | $N^*$ (100%) |
| 8 | 6.7% | RC (50%) + $N_T$ (50%) | 6.9% | RC (100%) |
| 9 | 7.7% | $N^*$ (100%) | 9.1% | RC (100%) |
| 10 | 3.2% | $N^*$ (100%) | 11.5% | RC (80%) + $N_T$ (20%) |
| 11 | 10.1% | $N^*$ (100%) | 13.5% | RC (75%) + $M$ (25%) |
| 12 | 14.3% | RC (55%) + $N_T$ (45%) | | |

**Fig. S7.**
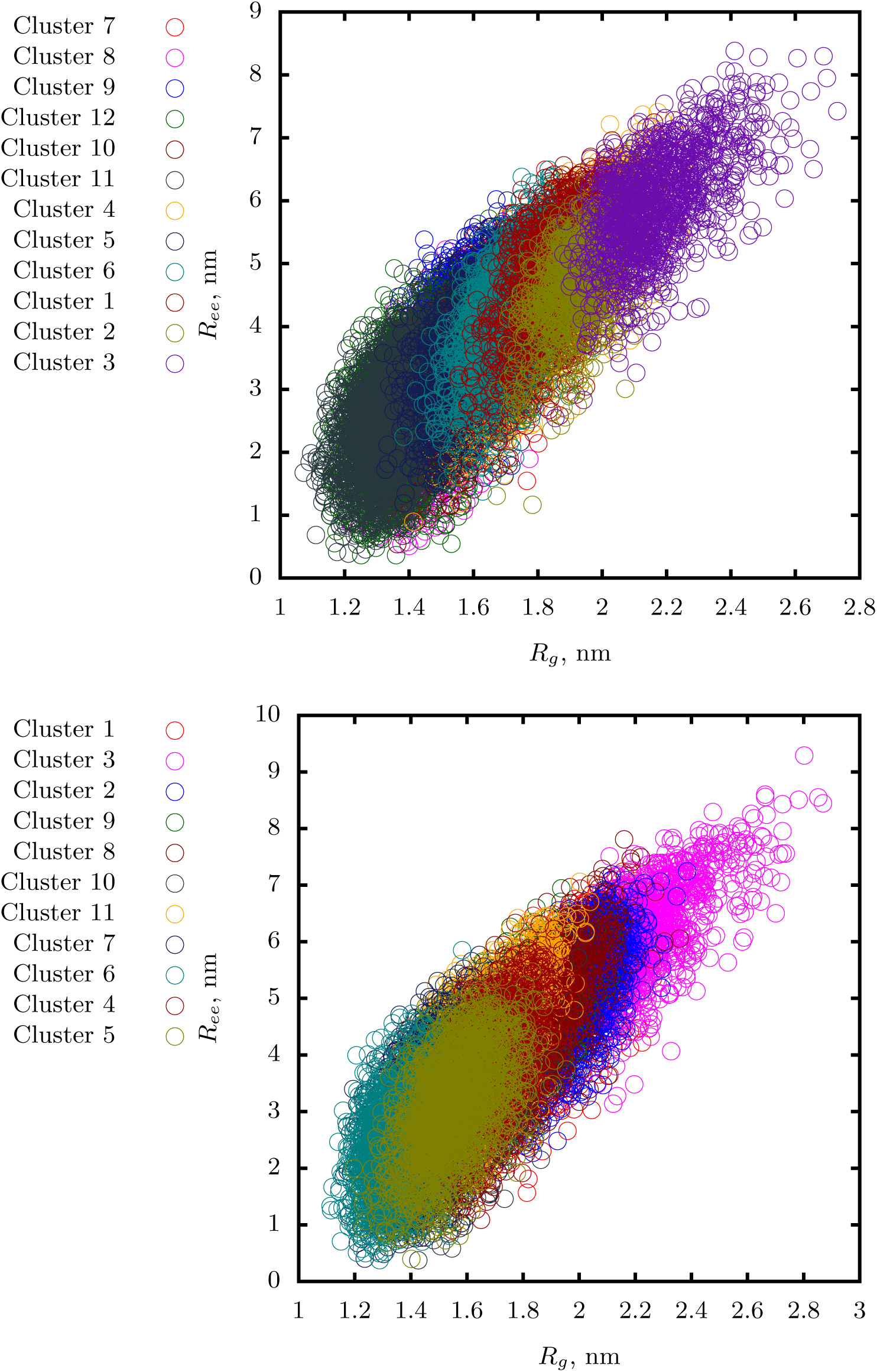
The different conformational clusters for A*β*40 (above) and A*β*42 (below) are well segregated in the landscape projected onto the *R*_*g*_ and *R*_*ee*_ coordinates, suggesting that the clustering scheme is robust.

**Fig. S8.**
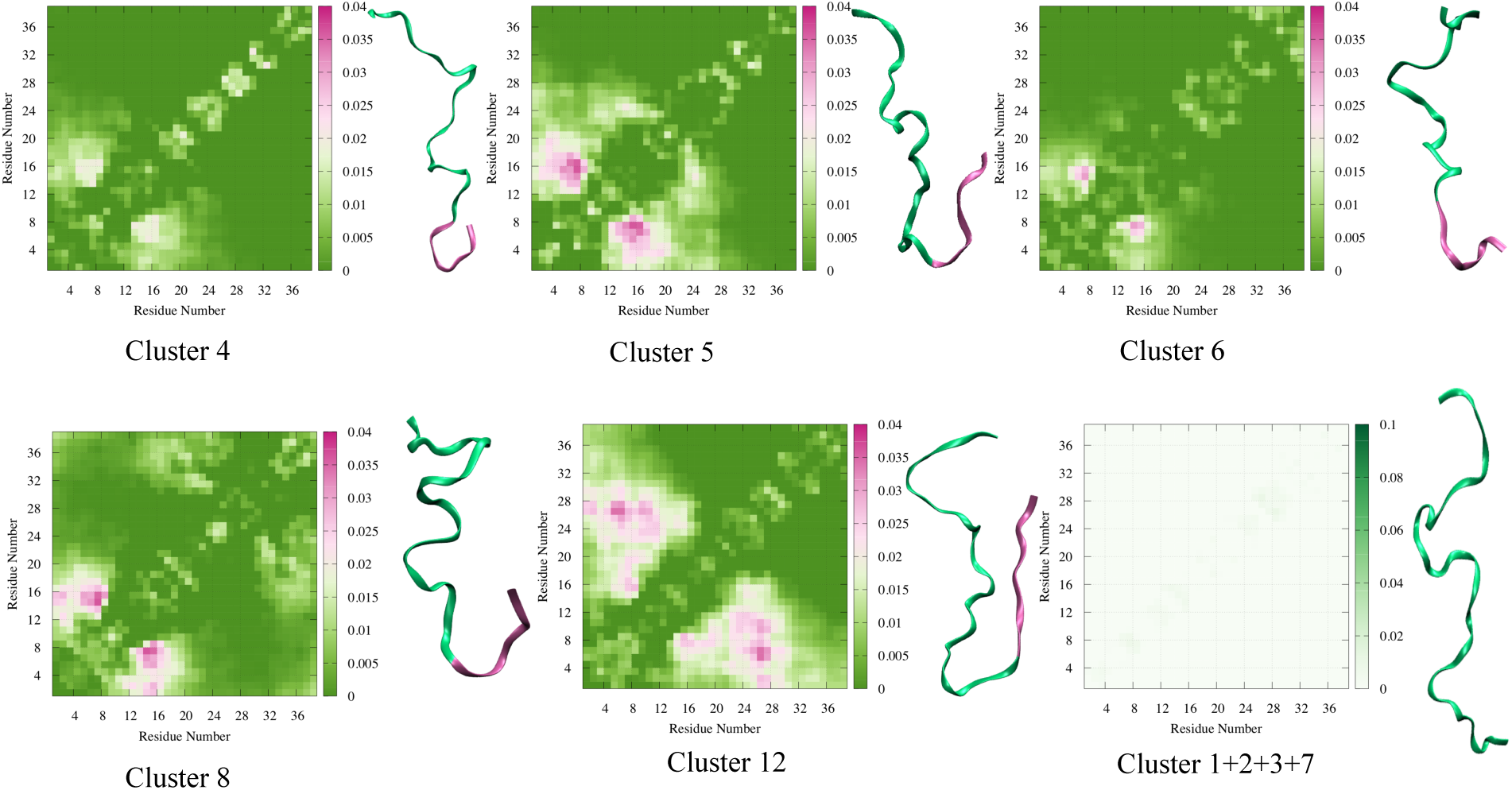
Clusters 4,5,6,8: Conformational clusters in A*β*40 with conformations with different extents of residual structuring near the N-terminus. Residues 1-10, which are disordered in the fibril structure, are shown in magenta, and residues 11-40, which are ordered in the fibril structure, are shown in light green. These clusters also consist of RC-like structures (see Table S3). The supercluster formed by clusters 1,2,3 and 7 consists of exclusively RC-like structures, and is characterized by a featureless contact map. Clusters in which the conformations with structural signatures of the fibril state are shown in the main text.

**Fig. S9.**
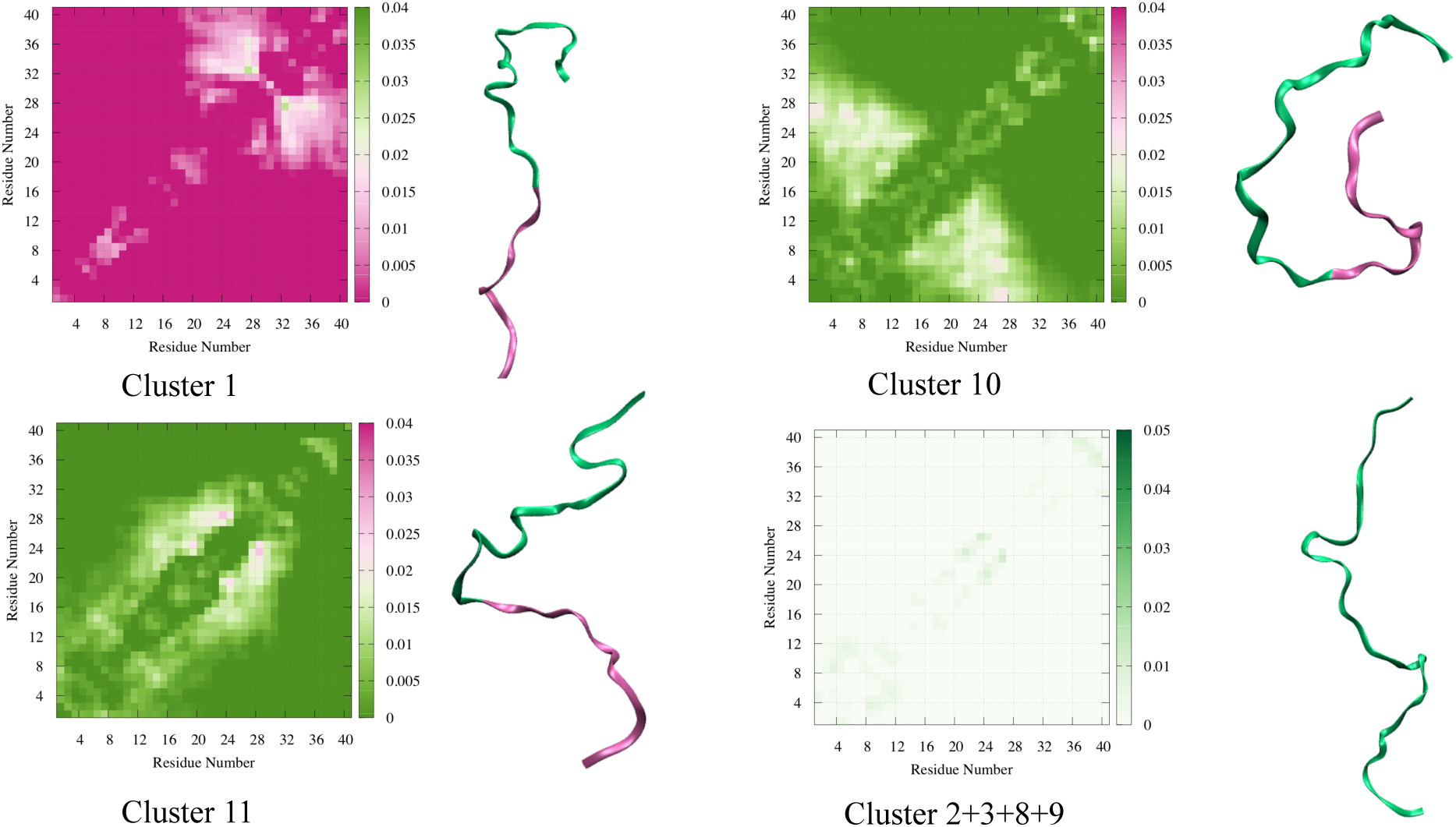
Different cluster families constituting the A*β*42 monomer ensemble. In cluster 1, the conformations have some structure in the C-terminus. Cluster 10 consists of structures with an ordered N-terminus. In cluster 11, both the termini are disordered. However, some residual structure is present near the middle of the peptide. In these clusters, residues 1-16, which form the disordered region in the fibril structure are shown in magenta, and residues 17-42, which form the ordered region of the fibril, are in green. These clusters also consist of RC-like conformations in addition to those exhibiting different extents of residual structuring. The supercluster formed by clusters 2,3,8 and 9 has a featureless contact map, and consists of exclusively RC-like conformations. The clusters which contain structures that have some resemblance to the monomer units of different A*β*42 polymorphs are shown in the main text.

**Fig. S10.**
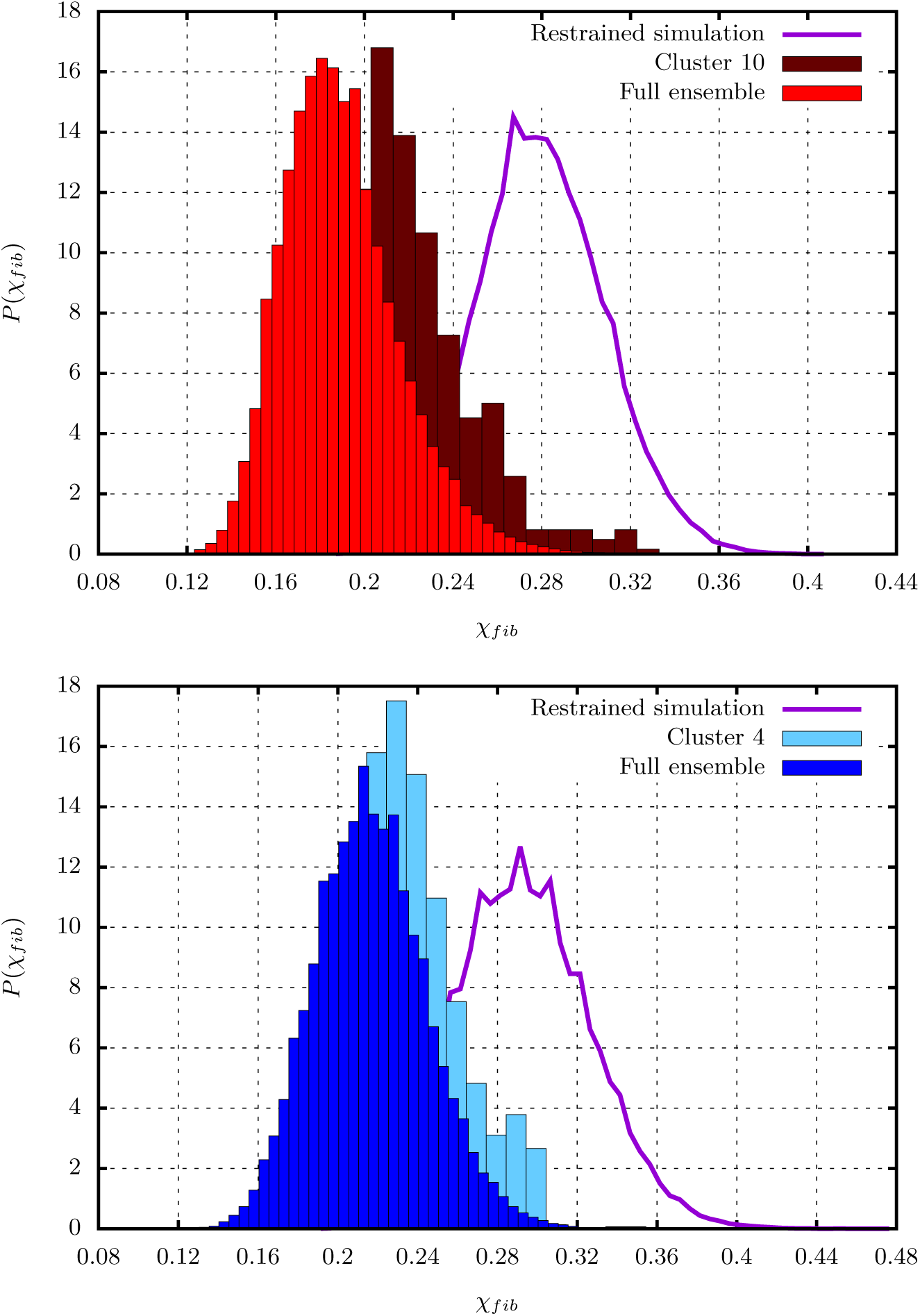
Top: The *χ*_*fib*_ distribution corresponding to cluster 10 from the A*β*40 conformational ensemble. This cluster exhibits structural signatures found in the fibril state. The distribution exhibits a minor peak near *χ*_*c*_=0.28, unlike the one computed from the full conformational ensemble. As is evident only a small population of structures within this cluster meet the strict geometric definition of a N* state. The purple curve denotes the distribution computed from restrained simulations. Bottom: The *χ*_*fib*_ distribution corresponding to cluster 4 from the conformational ensemble of A*β*42. This cluster exhibits signatures of the S-bend morphology. A minor peak appears at values near *χ*_*c*_indicating a subpopulation of structures present within this cluster could be geometrically characterized as N* configurations. The *χ*_*fib*_ distribution computed from restrained simulations is shown as a purple curve.

**Fig. S11.**
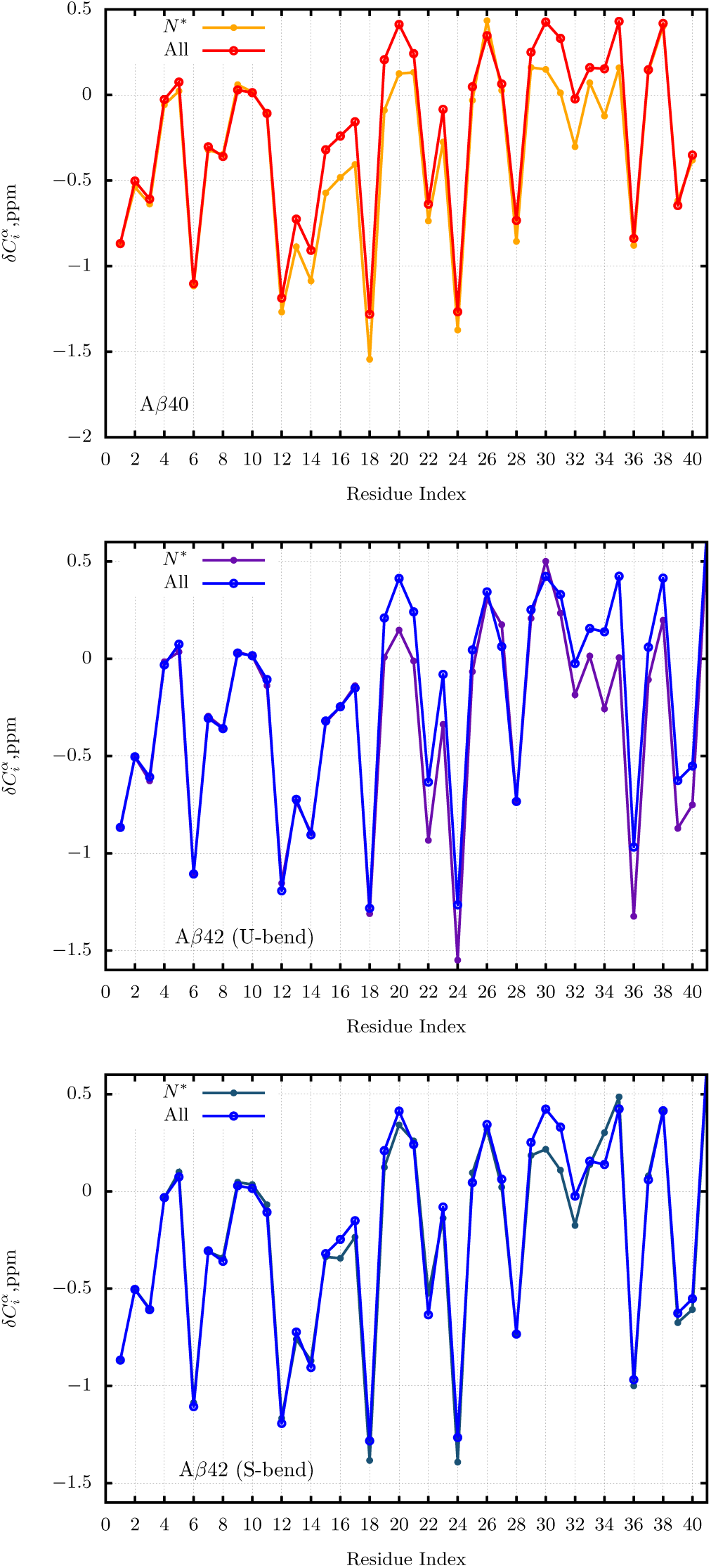
The residue-wise relative chemical shifts for the *N*\* ensemble overlaid on the profiles calculated using the full conformational ensemble. Top panel: *N*\* state of A*β*40, Middle panel: *N*\* state of A*β*42 having the U-bend topology, Bottom panel: *N*\* state of A*β*42 having the S-bend motif. The upshifted values of 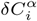 for some residues show that the *β* character is enhanced relative to the ensemble-average in the *N*\* state.

